# Rank-Preserving Alignment Enables Cross-platform Learning and Phenotyping for Single-Cell and Spatial Proteomics

**DOI:** 10.64898/2026.09.25.754510

**Authors:** Qunzhi Xu, Tian Le, Zhicheng Luo, Christopher Ly, Yinqiao Yan, Ye Zheng

## Abstract

Decades of antibody-based protein profiling have generated valuable cohorts across evolving cytometry, sequencing and spatial imaging platforms. Differences in marker coverage, signal scale and antibody performance hinder joint analysis and reuse of cohorts with long-term clinical follow-up. Widely used RNA-based integration methods prioritize shared embeddings rather than directly comparable protein measurements. To fill in the gap, we developed RAMP (Rank-preserving Alignment of Multi-platform Proteomics), an unsupervised framework that aligns shared proteins without cell-type annotations. It combines sample-specific anchoring and cross-platform cell matching with bounded monotone transformations. The corrected values retain each protein marker’s expression order across cells within a sample.

Across seven integration tasks, RAMP achieved the strongest overall balance of batch correction and cell-type preservation. RAMP also improved marker-threshold transfer while preventing expression-order reversals observed with some comparison methods. These reversals made cell types with originally high marker expression appear lower-expressing than other populations. Such overcorrection can mislead cell identification and protein comparisons, even when the integrated embedding shows improved dataset mixing. By preserving each marker’s expression order within a sample, RAMP protects the relationships needed to interpret normalized protein measurements. To investigate whether better harmonized inputs improve downstream learning, we used linear, nearest-neighbor and small neural-network models as a controlled, small-scale testbed. This approach tests an input quality question relevant to proteomic foundation models without undertaking large-scale pretraining. Under matched supervision, RAMP improved cross-platform annotation across all three model families and improved missing-marker prediction over unaligned inputs. These gains demonstrate the value of harmonized inputs for predictive learning and motivate evaluating RAMP as preprocessing for proteomic foundation models.

In bone marrow, we used high-quality single-cell protein measurements to impute the failed CODEX CD19 channel, recovering B-cell spatial distributions. The completed panel improved B-cell separation from natural killer cells in the joint embedding. In melanoma, a proliferating CD8 T-cell state near tumor cells was associated with longer survival. Transfer such a spatial information to CITE-seq connected tumor proliferation and infiltration state to an adhesion and activation program involving PD-1, LFA-1 and CD2. In lung cancer, we used single-cell CyTOF measurements to predict unmeasured HLA-ABC states in vascular cancer-associated fibroblasts profiled by spatial imaging mass cytometry (IMC). Higher predicted scores were associated with lower immune-cell fractions and worse disease-free survival in the IMC cohort. In head and neck cancer, transferring a fibroblast/macrophage spatial niche score to CITE-seq identified a CD39/TIGIT-associated CD8 T-cell phenotype. Protein and RNA profiles supported checkpoint-associated features, and transfer back to CODEX showed correspondence with measured LAG3. RAMP enables existing single-cell and spatial cohorts to support new discoveries by connecting molecular programs, tissue organization and clinical outcomes across platforms.

## Introduction

Single-cell protein profiling spans multiple technologies with distinct measurement principles. Mass cytometry (CyTOF) profiles dissociated cells at high throughput^1,2^; imaging mass cytometry (IMC), multiplexed ion beam imaging (MIBI) and cyclic fluorescence methods such as co-detection by indexing (CODEX) and multiplex immunofluorescence (mIF) preserve spatial context in intact tissue^3–6^; and sequencing-based assays quantify proteins using oligonucleotide-tagged antibodies; CITE-seq additionally measures transcripts in the same cells^7,8^. These technologies offer complementary views of cellular identity and tissue organization^9,10^. As research programs evolve, they accumulate data across instruments, antibody panels and assay generations. Even after a platform has been superseded, its measurements remain valuable, particularly when linked to long-term clinical follow-up. Reusing these data requires a way to compare protein measurements across technologies and connect information available in one dataset to information missing from another.

This comparison is complicated by differences in antibody clones, conjugates and detection chemistry, which alter the recorded signal for the same protein. The Blueprint PD-L1 studies, for example, found comparable tumor-cell staining with several assays but systematically lower sensitivity with SP142^11,12^. Antibody performance also varies with reagent lot and application^13,14^. Variability persists within a single technology: the multisite MITRE study reported average intersite cell-density agreement of *R*^2^ = 0.72–0.75 for a standardized multiplexed immunofluorescence workflow^15^. Across imaging platforms, similar population-level estimates can coexist with marker-dependent differences at single-cell resolution^16^. Such differences complicate the transfer of analysis pipelines and biological interpretations between datasets^17^.

The requirements for harmonization follow from how protein measurements are used. Many downstream analyses operate directly on marker values. Flow and mass cytometry define populations through gating, and clinical workflows rely on standardized immunophenotyping protocols and reproducible population estimates^18,19^. Efforts such as EuroFlow therefore standardize instrument settings and antibody panels to make measurements comparable across instruments and sites^20,21^. Spatial proteomics likewise assigns phenotypes through manual thresholds or models that encode marker positivity^22–24^, yet variation in signal intensity makes universal cutoffs difficult to apply^25^. The intensity scale can change the resulting biological interpretation: in multiplexed imaging, normalization choices can affect cell-type assignments more strongly than clustering choices^26^. Marker-level quality control also depends on examining the measured signals to identify staining failures and other artifacts^27,28^.

Integration methods differ in how well their outputs support these analyses. Many widely used approaches focus on aligning cell relationships or constructing shared representations^29–31^. Although these outputs support clustering, visualization and cell matching, an aligned embedding alone does not provide a corrected CD8 measurement to which a gate can be applied. Marker-level analyses may therefore continue to use the original, unaligned measurements. The dependence of downstream analyses on the type of integration output is recognized in transcriptomics^32^; in proteomics, it directly affects workflows built around individual marker values and thresholds.

Methods such as cyCombine, CytoNorm and CytofRUV return corrected marker values^33–35^, but correction in expression space does not necessarily preserve the ordering of cells within a sample. Even a monotone transformation applied separately to different cell clusters can change the relative order of cells across clusters. This distinction matters because correction can alter the marker relationships used to define cell populations. Evaluations in single-cell transcriptomics have shown that batch correction can modify data even when no batch effect is present^36^, and that commonly used integration metrics can miss overcorrection^37^. Earlier proteomic studies also illustrate how technical artifacts can produce misleading biological conclusions^38^. Monotone transformation has been shown to be valuable for aligning antibody-derived tag measurements across CITE-seq studies^39^. Extending this principle across measurement technologies requires reconciling different signal scales while retaining the within-sample marker relationships used in downstream analysis.

Here we present RAMP (Rank-Preserving Alignment of Multi-Platform Protein Data), an unsupervised framework that aligns protein measurements across two or three platforms without cell-type annotations. RAMP anchors each marker’s low-expression population within each sample, establishes cross-platform cell correspondences using complementary representations, and fits sample- and marker-specific monotone transformations with bounded displacement. The result is a shared normalized intensity scale that preserves within-sample cell ordering. We evaluate RAMP on 17 datasets spanning seven platforms, 299 sample units, four cancer types and healthy bone marrow. RAMP ranks first in five of seven integration tasks; across the four original benchmarks, it improves the mean balanced accuracy of fixed-threshold transfer from 0.648 to 0.869. Under matched supervision, RAMP improved cross-platform annotation across three model families and increased missing-marker prediction performance relative to unaligned inputs. We further use the aligned measurements for downstream prediction and state transfer: combining RAMP preprocessing with reference-based completion of a failed CD19 channel in 12 CODEX bone-marrow donors, transferring a spatially defined CD8 T-cell state into dissociated CITE-seq data, predicting an unmeasured HLA-ABC state in a lung cancer cohort with 280 patients in the disease-free survival analysis, and identifying proteins associated with a spatial niche in independently profiled single cells. These applications connect protein states, tissue organization and clinical outcomes across datasets that were collected separately.

## Results

### RAMP aligns protein measurements with bounded monotone transformations

RAMP aligns the shared proteins of two or three datasets without requiring cell-type annotations (Fig. 1A). It first applies an adaptive arcsinh transformation within each sample and marker, identifies a low-expression population by one-dimensional *k*-means, and anchors and scales the resulting distribution. It then establishes cross-platform cell correspondences from complementary source-scale and preprocessed representations. Bidirectional matching support and, for three datasets, cycle consistency inform these correspondences. Finally, matched cells provide targets for sample- and marker-specific monotone transformations, with calibrated correction strength and a fixed displacement ceiling.

**Figure 1:**
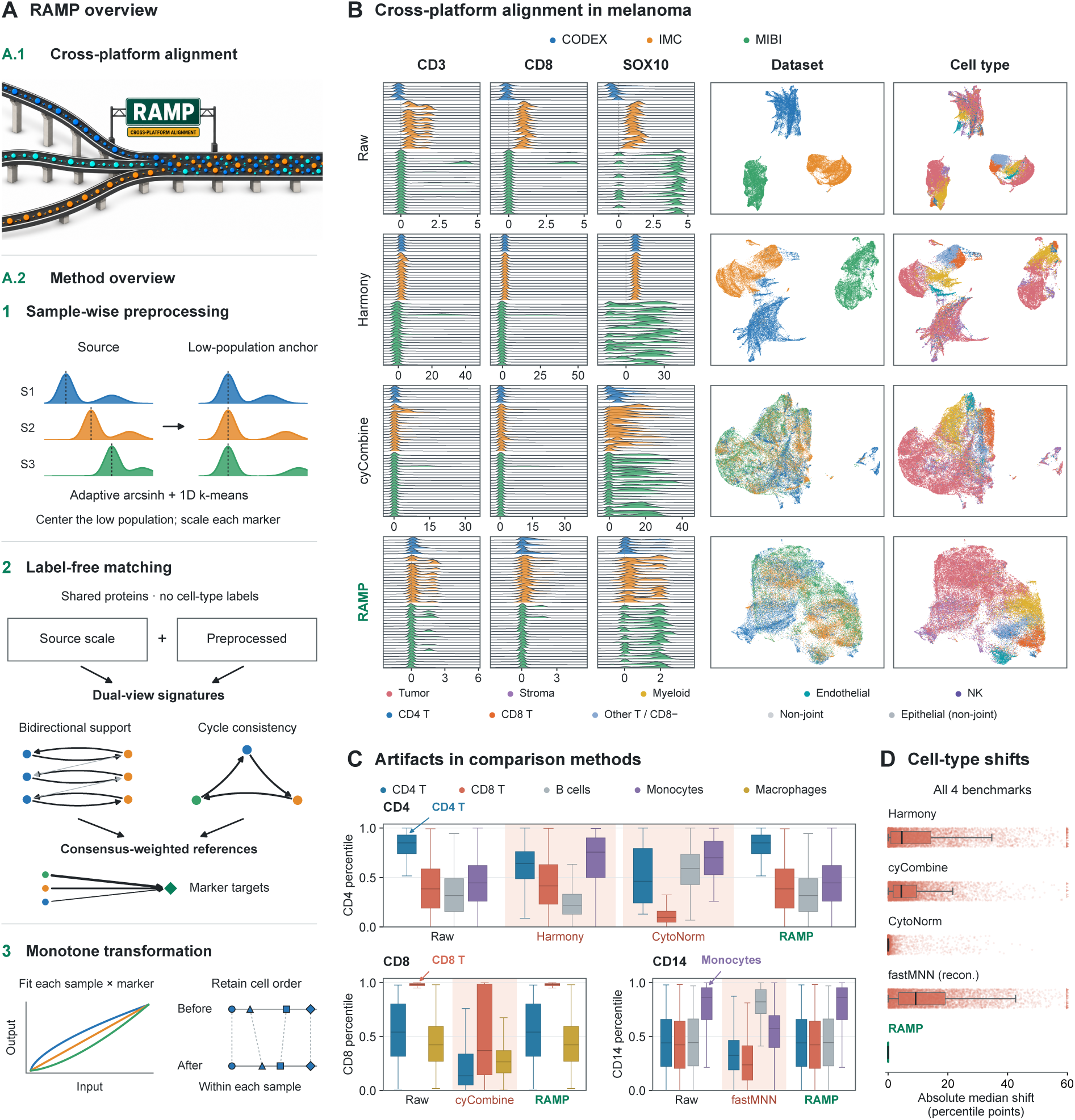
RAMP aligns protein measurements while preserving within-sample marker order. **(A)** Concept and algorithm: low-expression anchoring, label-free cross-platform matching and bounded monotone transformation. Diagrams are schematic. **(B)** Alignment of 60,000 melanoma cells using 12 shared proteins. CODEX, IMC and MIBI denote Hickey et al. ^40^, Hoch et al. ^41^ and Vesely and Johnson ^42^, respectively. Ridges of CD3, CD8 and SOX10 across 46 sample units accompany identical UMAP coordinates colored by dataset and cell type. Ridges are peak-normalized, with method-specific display windows (Methods). **(C)** Selected marker-percentile changes in CyTOF CD4, CODEX CD8 and CyTOF CD14. Percentiles use all cells within each sample. Exploratory examples illustrate population-specific changes; discrimination AUROCs are in Results. **(D)** Absolute change in cell-type median marker percentile across 6,173 sample × cell type × marker combinations from 103 sample units in four benchmarks, using identical combinations and Raw references for Harmony, cyCombine, CytoNorm, reconstructed fastMNN and RAMP. Boundary triangles mark values beyond 60; box statistics use untruncated data. FastMNN reconstruction is diagnostic; MaxFuse has no corresponding marker matrix. Cell-type labels are used for evaluation and display, not alignment. Boxes show quartiles and whiskers extend to the most extreme values within 1.5 interquartile ranges.

The output is a cell-by-marker matrix on a shared, normalized intensity scale. For any two cells in the same sample, a higher input value for a marker cannot become a lower output value for that marker. This order-preservation property follows from the non-decreasing transformation, including the bounded update; ties may be introduced, but strict reversals are excluded (Methods). The final correction is limited to 0.75 units relative to the preprocessed value. Matching determines the proposed calibration, while monotonicity and the displacement bound constrain how it can modify the measurements.

We first evaluated RAMP on four frozen three-platform benchmarks comprising 270,732 cells in 104 sample units from bone marrow, breast cancer, melanoma and lung cancer^40–51^. In the three-platform melanoma benchmark, containing 60,000 cells and 12 shared proteins, source datasets differed in their marker distributions and separated in UMAP space. RAMP brought the distributions onto a common scale and increased dataset mixing while retaining organized cell-type structure (Fig. 1B).

### Marker-level audits identify population-specific changes in comparison outputs

To examine properties that an integrated embedding does not directly measure, we compared marker distributions before and after correction. Values were converted to within-sample percentiles, allowing the relative positions of annotated cell populations to be compared across methods (Fig. 1C). In one bone-marrow CyTOF sample, CD4 separated CD4 T cells from monocytes with AUROC 0.926 before correction. AUROC fell to 0.378 with Harmony and 0.322 with CytoNorm, reversing the relative ordering of these populations. CytoNorm also changed the CD4-T-versus-B-cell AUROC from 0.968 to 0.403. In a breast CODEX sample, cyCombine reduced the CD8-T-versus-macrophage AUROC from 0.976 to 0.610 and shifted the median CD8 T-cell percentile from 0.983 to 0.371. In a second bone-marrow sample, reconstructed PCA-fastMNN marker coordinates changed the monocyte-versus-B-cell CD14 AUROC from 0.811 to 0.149. RAMP retained the source percentile profiles in these examples.

These observations illustrate the distinction between correction of a multivariate representation and preservation of an individual marker axis. CytoNorm fits cluster-specific monotone splines, whereas cyCombine applies ComBat within self-organizing-map nodes^33,34^. Different transformations across partitions can change the ordering of cells assigned to different groups. Harmony operates directly in marker coordinates, but its cell-dependent corrections do not impose marker-wise monotonicity. FastMNN’s primary output is a corrected embedding; the reconstructed marker coordinates examined here are a diagnostic representation rather than calibrated protein abundance. Cell- or cluster-dependent corrections can therefore change the ordering of cells along an individual marker. RAMP constrains each sample–marker distribution to a common monotone transformation, excluding strict within-sample order reversals by construction.

We then quantified distribution shifts across all four benchmarks. Among 6,173 eligible sample × cell type × marker combinations from 103 sample units, each tested marker-valued comparison output had a tail of large absolute changes in cell-type median percentile (Fig. 1D). RAMP’s median change was zero and its maximum was 0.0821 percentile points, attributable to changes in ties. Across these examples, correction altered marker relationships used to distinguish established cell populations. The broader audit showed that RAMP retained population positions within sample-level marker distributions. These results demonstrate the value of preserving marker ordering alongside improving cross-platform integration.

### RAMP achieves the best overall rank across seven integration tasks

We compared RAMP, MaxFuse, fastMNN, cyCombine, CytoNorm and Harmony on the four benchmarks and three additional tasks drawn from the lung, melanoma and head-and-neck applications below. All methods used the same cells within each task; MaxFuse additionally used the available full protein panels in its active feature space (Methods). The seven tasks together covered 17 source datasets, seven technologies and 299 distinct dataset-specific sample units. Some tasks shared source cohorts, and sample units were not paired across platforms.

RAMP had the highest seven-task mean for Louvain adjusted Rand index against harmonized cell types and for macro-averaged cell-type silhouette in UMAP space (Fig. 2A,B). Across the main 23-metric summary, RAMP had the highest mean relative performance in cell-type structure, platform and donor mixing, and UMAP-based separation; fastMNN had the highest mean relative performance in cross-platform label transfer (Fig. 2C; Supplementary Figs. S8 and S9). Categories were weighted equally so that the 14 mixing metrics did not dominate the summary. The resulting overall mean rank was 2.01 for RAMP, compared with MaxFuse (2.70), fastMNN (2.89), cyCombine (3.79), CytoNorm (4.39) and Harmony (5.23; Fig. 2D). RAMP ranked first in five tasks and second in two tasks.

**Figure 2:**
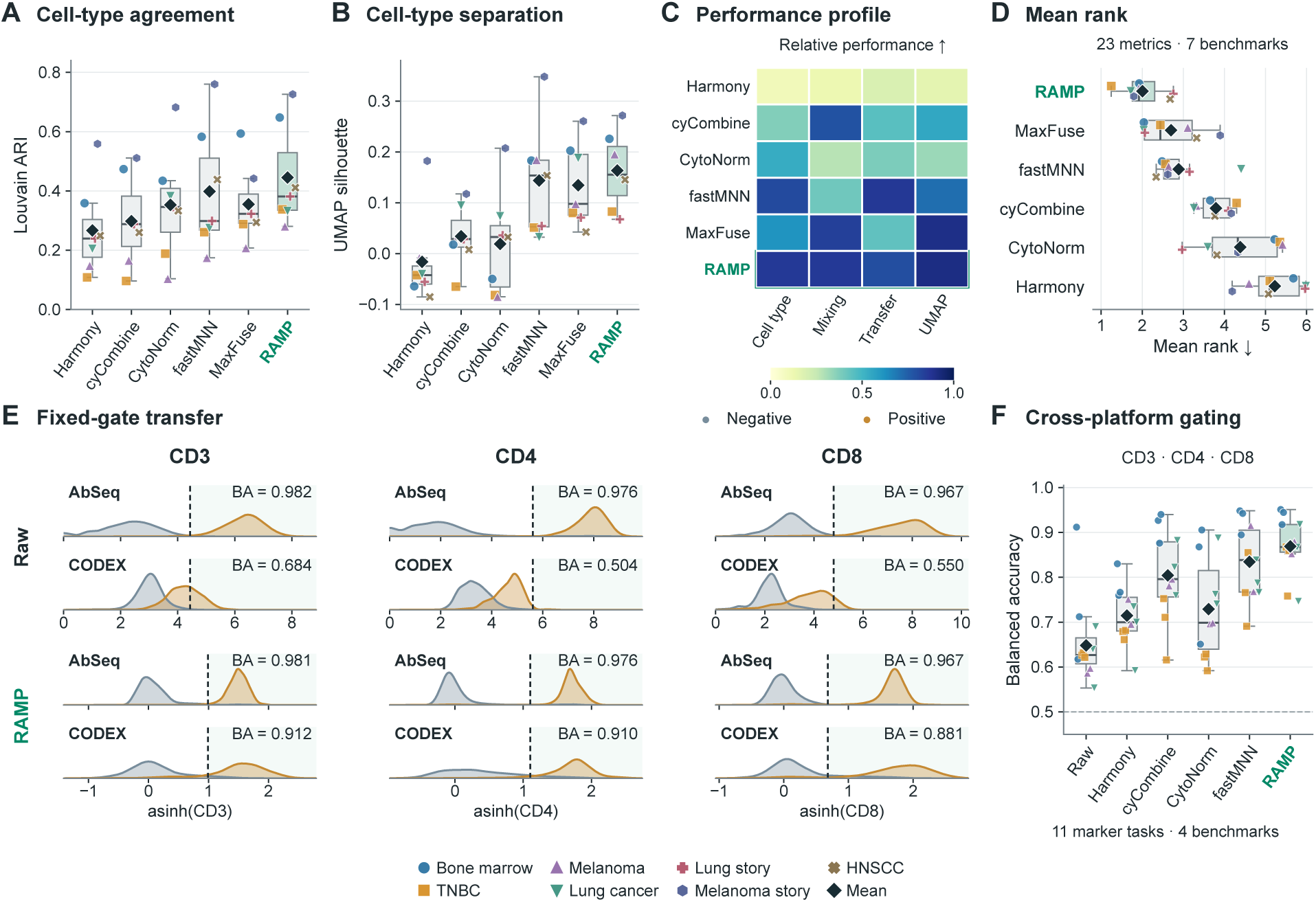
Integration performance and cross-platform portability of marker thresholds. **(A,B)** Louvain adjusted Rand index against harmonized cell types and macro-averaged cell-type silhouette in UMAP space. Each point is one of seven integration tasks, averaged over 20 matched resamples of 5,000 cells per source dataset. Black diamonds show the task mean; boxes and whiskers use the same conventions as the overview (Fig. 1). **(C)** Relative performance over 23 metrics in four categories: cell-type structure (5), platform and donor mixing (14), cross-platform label transfer (2) and UMAP (2). Direction-adjusted scores are scaled between the worst and best methods within each task and metric, then averaged within category and equally across tasks. **(D)** Mean method rank, giving equal weight to the four categories. Points represent tasks and diamonds their mean. RAMP ranks first in five tasks and second in two tasks, with an overall mean rank of 2.01. The main metric subset was chosen after result inspection; full 31-metric and alternative-weighting summaries are reported in Methods and the supplementary CSV data. **(E)** Bone-marrow CD3, CD4 and CD8 gates calibrated on three AbSeq donors and transferred to 12 CODEX donors. Rows show Raw AbSeq, Raw CODEX, RAMP AbSeq and RAMP CODEX. Gold and gray-blue indicate reference-positive and negative populations. Each method’s source-calibrated gate (dashed line) is applied unchanged to its target values. Target labels do not select gates. BA, donor-averaged balanced accuracy. **(F)** Gate transfer across four benchmarks and 11 marker tasks. Each point averages six platform-transfer directions; diamonds average tasks within benchmark, then benchmarks equally. Overall BA is 0.648 for Raw, 0.869 for RAMP and 0.835 for reconstructed fastMNN. A–D use fastMNN’s corrected components; F uses diagnostic reconstructed marker coordinates. MaxFuse has no corrected marker matrix for this test. Gate-transfer evaluations use frozen jointly aligned cohorts and do not enter the integration rank.

RAMP retained the best overall rank across all four sensitivity summaries, combining either the 23 primary or all 31 evaluated metrics with equal-category or equal-metric weighting (Methods). Its leading overall performance therefore persisted across both metric sets and aggregation schemes. These sensitivity analyses accompany the primary reporting set, which was selected after inspection of the results.

### Aligned marker values improve cross-platform gate transfer

We next tested whether the corrected values support a downstream operation that depends directly on marker scale: applying a positivity threshold learned on one platform to another. For each method and marker, we calibrated a threshold using source-platform labels alone and applied that threshold unchanged to the target platform. Thus, the test measures agreement between source and target scales after correction, separately from order preservation.

For bone-marrow CD3, CD4 and CD8, thresholds were calibrated on three AbSeq donors and evaluated on 12 CODEX donors (Fig. 2E). Without alignment, target balanced accuracy was 0.684, 0.504 and 0.550, respectively. With RAMP, the corresponding accuracies were 0.912, 0.910 and 0.881. Across 11 marker tasks and six transfer directions per task in the four benchmarks, the equally weighted benchmark mean increased from 0.648 for Raw to 0.869 for RAMP. The next highest mean was 0.835 for reconstructed fastMNN coordinates (Fig. 2F). MaxFuse was excluded from this value-level test because the evaluated implementation returned an embedding and cell matches without a corrected marker matrix. The gate-transfer results show that RAMP’s corrected values improve the portability of marker-level decisions across the tested platforms.

### RAMP supports cross-platform predictive learning

We next asked whether alignment makes measurements from different platforms more useful for training a shared prediction model. We evaluated cell-type annotation and missing-marker prediction using linear models, *k*-nearest neighbors and a single-hidden-layer neural network (Fig. 3A). RAMP was compared with five alignment methods, an assay-ready unaligned baseline, and three simple preprocessing strategies: upper-clipped *z*-score, min–max scaling and within-sample percentile transformation. The methods used the same frozen cells and shared-marker information. Prediction models were trained and selected using source-cohort supervision; target answers were reserved for evaluation. The unlabeled target expression was included in alignment, so this experiment evaluates prediction within jointly processed cohorts rather than alignment of a previously unseen cohort (Methods).

**Figure 3:**
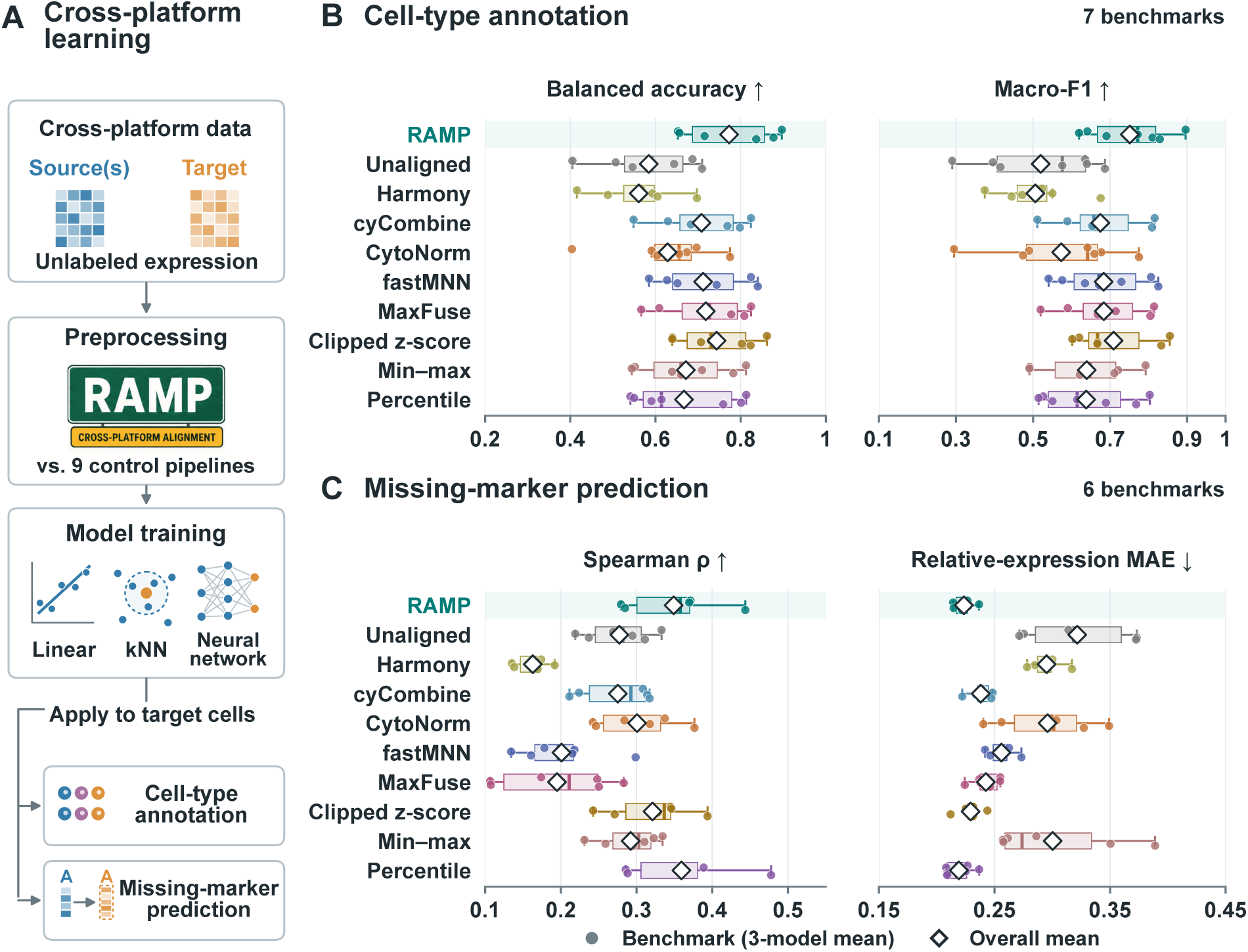
RAMP supports cross-platform prediction across model families. **(A)** Evaluation workflow. Unlabeled source and target expression is processed by RAMP or nine control pipelines. Linear, *k*-nearest-neighbor and neural-network models use source supervision for training and model selection, then predict target cell types or withheld-marker expression. For marker prediction, the marker is removed from all platforms before preprocessing and alignment are refitted. Diagrams are schematic. **(B)** Cell-type annotation across seven benchmarks and 34 directed transfers, using 500 source training and 512 validation cells. Training and scoring use the updated task-specific class scopes listed in Methods; these scopes were revised after exploratory review of the original benchmark. **(C)** Missing-marker prediction across six benchmarks and 79 benchmark–marker combinations. Spearman correlation is computed within sample units; mean absolute error (MAE) compares predictions with a common 0–1 relative-expression target, not absolute protein abundance. Source calibration uses an additional 64 cells per unit (Methods). The lung application benchmark uses the QC-filtered Senosain CyTOF reference described in Methods. No annotation-class filter is applied to this regression task. In B,C, each point is a benchmark score averaged equally across the three model families. Open diamonds show benchmark-equal means; boxes show quartiles and the median, and whiskers extend to the most extreme values within 1.5 interquartile ranges. All benchmark points are displayed. Three random seeds were used, with source training and validation support checked before comparison. Benchmark configurations share some source cohorts; points are descriptive summaries, not independent-patient replicates.

For cell-type annotation, we evaluated 34 directed transfers across seven benchmark configurations, using 500 source training cells and 512 source validation cells. In the updated, task-specific classification scopes (Methods), RAMP achieved mean balanced accuracy of 0.773 and macro-F1 of 0.751, compared with 0.584 and 0.520 without alignment (Fig. 3B; Supplementary Figs. S10 and S11). The next highest means were 0.744 and 0.709 for clipped *z*-score preprocessing. With identical eligible labels, evaluation cells and source-supervision budgets across methods, RAMP ranked first for both metrics in six of seven benchmarks and second in HNSCC. It also achieved the highest benchmark-mean scores within each of the three model families, showing that the annotation benefit extended across linear, nearest-neighbor and neural-network classifiers.

For missing-marker prediction, each marker was removed from every platform’s inputs before rebuilding preprocessing and alignment. Models learned the relationship between the remaining markers and withheld source measurements, then predicted the same marker in the target cohort. The evaluation covered all 79 eligible benchmark–marker combinations in six benchmarks (Fig. 3C; Supplementary Figs. S12 and S13). RAMP achieved the highest mean within-unit Spearman correlation and lowest relative-expression MAE among the six alignment methods. Compared with unaligned inputs, correlation increased from 0.277 to 0.349 and MAE decreased from 0.322 to 0.223. Across all ten processing pipelines, RAMP ranked second for both endpoints, following percentile preprocessing (Spearman correlation 0.359; MAE 0.219). These results show that RAMP-supported prediction recovered withheld-marker variation across platforms while improving accuracy over the other tested alignment pipelines.

### Reference-based completion recovers B-cell-associated CD19 signal

We investigated whether measurements from reference platforms could help recover a failed channel in a spatial dataset. The healthy bone-marrow cohort contained 88,031 cells from CODEX, CyTOF and AbSeq^46–48^. In CODEX, acquired DAPI resolved cells in a field with little detectable CD19 signal (Fig. 4A). A label-free anomaly screen, combining reference-based prediction disagreement with distribution diagnostics, ranked CD19 as the most anomalous channel in all 12 CODEX donors, allowing ties. Consistently, observed CD19 separated annotated B cells from CD4/CD8 T cells with median within-donor AUROC 0.53 in CODEX, compared with 1.00 in CyTOF and 0.99 in AbSeq (Fig. 4B).

**Figure 4:**
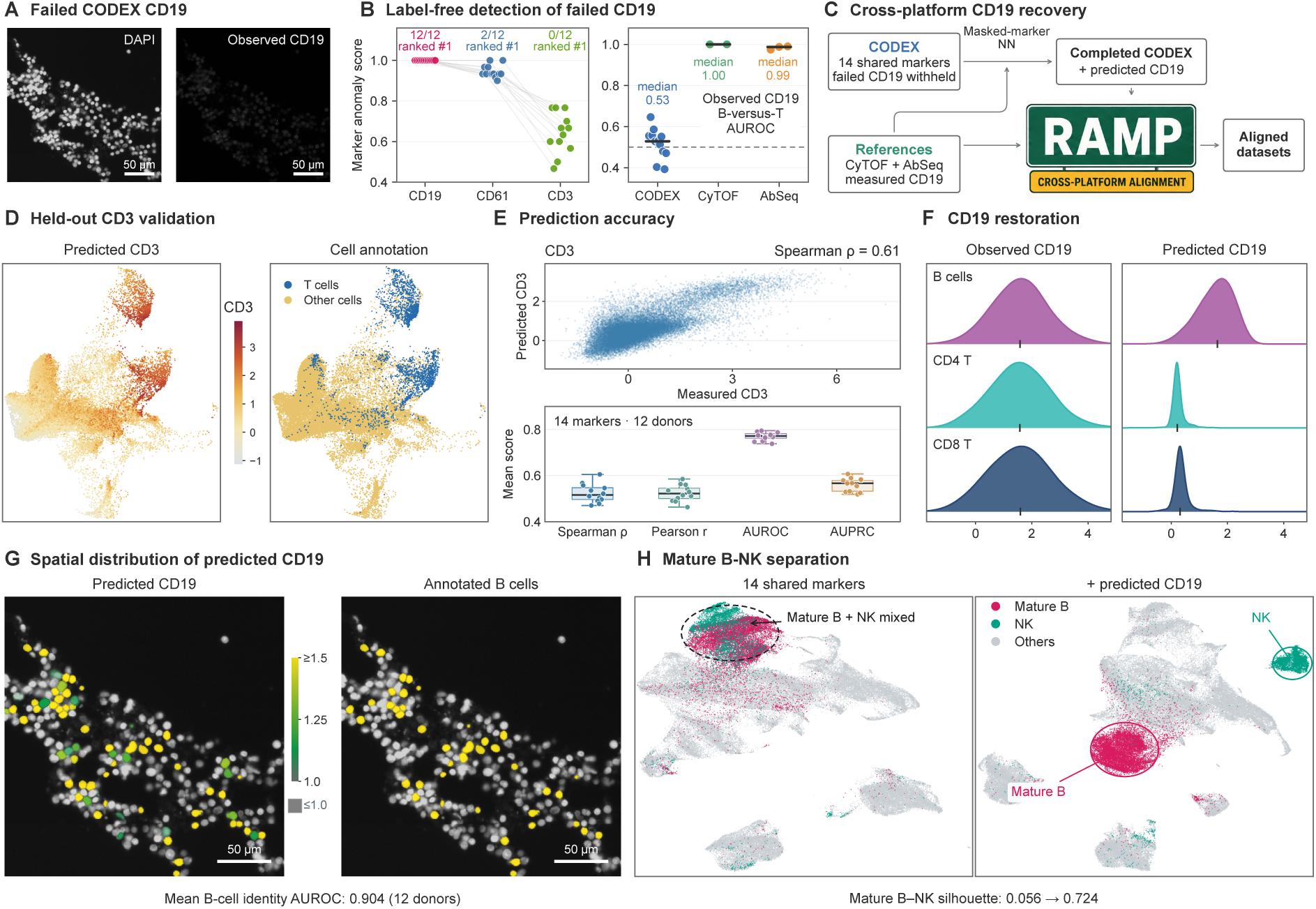
Reference-based completion of a failed CODEX CD19 channel in bone marrow. The cohort contains 30,000 CODEX cells from 12 donors, 30,000 CyTOF cells from two donors and 28,031 AbSeq cells from three donors ^46–48^. **(A)** Acquired DAPI and CD19 images from the same registered H14 field shown in G. Scale bars, 50 µm. **(B)** Label-free channel anomaly scores (left) and observed CD19 B-versus-CD4/CD8-T-cell AUROC (right). Each point is a donor. CD19 has the highest anomaly rank in all 12 CODEX donors, allowing ties. Dashed line, AUROC 0.5. **(C)** Workflow: low-expression preprocessing of 14 functioning shared markers, masked-marker neural-network completion using reference CD19 targets, and RAMP alignment of the completed panel. Failed CODEX CD19 is excluded from inputs and targets. **(D)** Permanent-holdout CD3 validation. Both plots show 30,000 CODEX cells on a joint 88,031-cell UMAP computed without CD3 or CD19. Left, predicted CD3; right, annotated CD4/CD8 T cells versus other cells. CD3 is excluded from all predictor inputs and CODEX targets but retained as a reference target. **(E)** Measured versus predicted CD3 for all sampled CODEX cells (Spearman *ρ* = 0.615; top). Bottom, validation of all 14 functioning markers: each point is one donor’s mean across marker-specific tasks. Median donor means are AUROC 0.772, average precision 0.567, Spearman 0.516 and Pearson 0.522. These are marker holdouts, not donor holdouts. **(F)** Observed and predicted CD19 distributions in B, CD4 T and CD8 T cells. Densities are independently peak-normalized; ticks indicate medians. **(G)** Continuous predicted CD19 and annotated B cells over acquired DAPI and provider cell masks. The fixed predictor was applied to all 83,384 H14 atlas cells; this selected field contains 251 cells, including 43 B cells (ROI selection in Methods). Gray denotes values at or below the display lower limit of 1.0 and yellow saturation begins at 1.5; neither is a biological positivity threshold. The footer reports mean B-versus-non-B AUROC across 12 sampled donors (0.904), not a field-specific metric. **(H)** Joint alignments without and with completed CD19, displaying mature B, NK and other cells. B-versus-NK UMAP silhouette increases from 0.056 to 0.724. Cell labels and spatial coordinates are excluded from predictor and alignment fitting.

We first applied RAMP’s sample-specific low-expression preprocessing to the 14 normally measured shared markers. A masked-marker neural network then learned marker relationships from the three platforms, using measured CD19 targets only from CyTOF and AbSeq. Failed CODEX CD19 values entered neither the inputs nor the targets. The two reference-platform predictions were averaged to complete the CODEX panel, which subsequently underwent joint RAMP alignment (Fig. 4C). This separates reference-based marker completion from the final alignment of the completed panel.

To evaluate prediction where an observed target was available, we permanently withheld each of the 14 functioning markers from every platform’s inputs and from CODEX training targets, then trained a separate model for each task. Predicted CD3 localized to annotated T cells in the joint UMAP (Fig. 4D) and correlated with measured CD3 across all 30,000 sampled CODEX cells (Spearman *ρ* = 0.615). Across donors, the median of donor-level means over the 14 tasks was 0.772 for AUROC, 0.567 for average precision, 0.516 for Spearman correlation and 0.522 for Pearson correlation (Fig. 4E). These marker-holdout results support the use of reference-platform measurements to recover cell-level expression patterns in CODEX.

Predicted CD19 was higher in B cells than in CD4 or CD8 T cells, whereas the observed channel showed strongly overlapping distributions (Fig. 4F). Applying the fixed predictor to the full H14 atlas allowed predictions to be overlaid on acquired DAPI using the original cell masks. High predicted values corresponded spatially to annotated B cells (Fig. 4G); across the 12 sampled donors, mean B-versus-non-B identity AUROC was 0.904. This evaluates correspondence with B-cell identity and tissue distribution, rather than accuracy against an intact CD19 channel. Finally, including completed CD19 in the joint alignment increased the UMAP silhouette between mature B and NK cells from 0.056 to 0.724 (Fig. 4H). Together, reference-based completion recovered B-cell-associated CD19 signal in a failed spatial channel and improved mature-B-versus-NK separation in the joint embedding. This application illustrates how RAMP preprocessing and subsequent alignment can support the recovery and downstream use of information lost through channel failure.

### A spatial CD8 T-cell state transfers to dissociated melanoma cells

We next asked whether a protein correlate of a spatially defined state could be recognized in dissociated cells. In pretreatment melanoma IMC^52^, we defined the target as antigen-experienced CD8 T cells with Ki67 positivity probability ≥ 0.5 and a melanoma-cell centroid within 30 µm (Fig. 5A). Among 30 patients receiving immune checkpoint therapy, a higher measured target-cell fraction was associated with longer overall survival: median survival was 554 versus 158 days after a median split (log-rank *P* = 0.000157; Fig. 5B). This association motivated transfer of the observed state.

**Figure 5:**
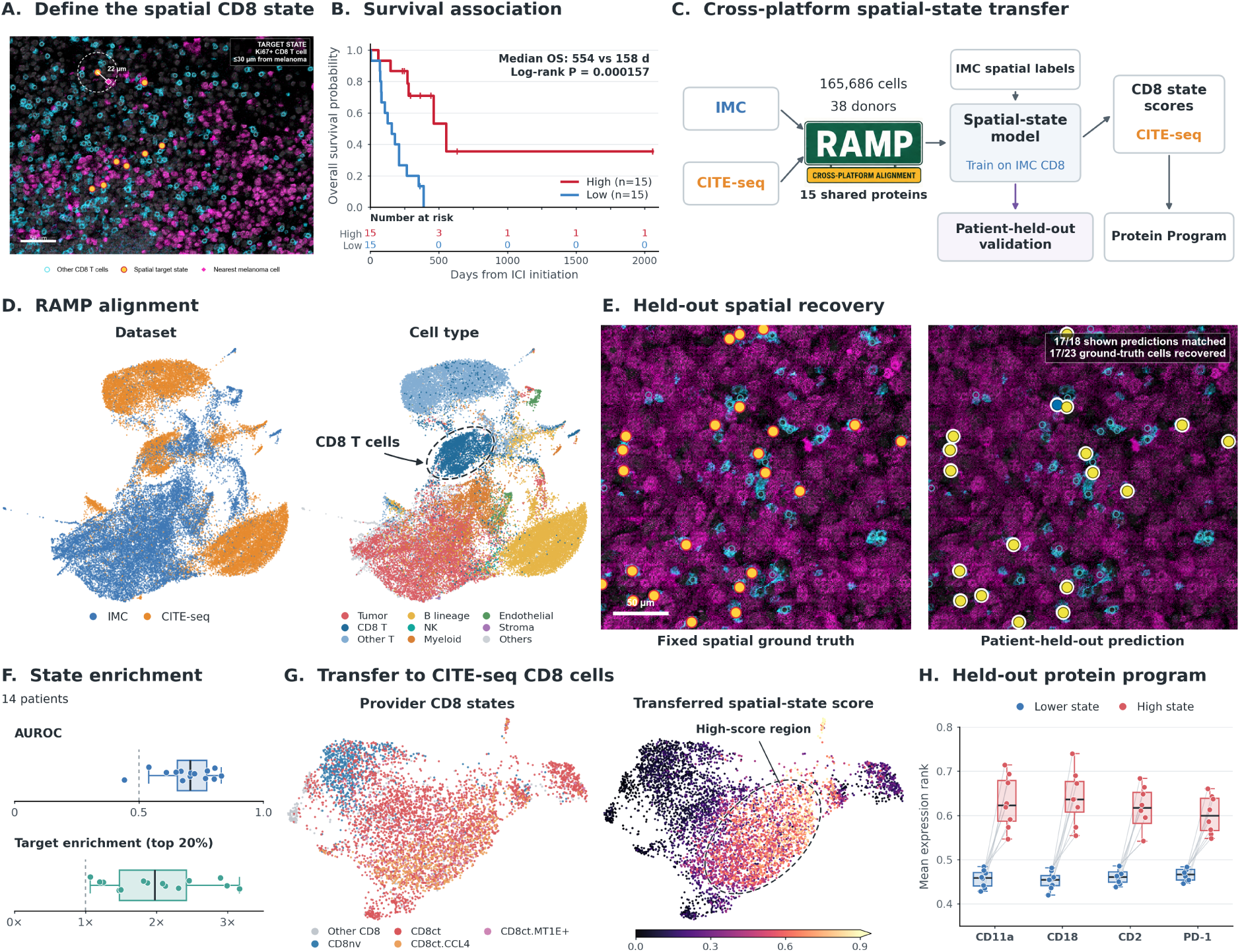
Transfer of a survival-associated spatial CD8 T-cell state from melanoma IMC to CITE-seq. The analysis combines pretreatment IMC from Moldoveanu et al. ^52^ with stage III melanoma-positive sentinel-node CITE-seq from Engelbrecht et al. ^53^. **(A)** The source target comprises provider-annotated antigen-experienced CD8 T cells with Ki67 positivity probability ≥ 0.5 and nearest melanoma-cell centroid within 30 µm. **(B)** Overall survival after checkpoint-therapy initiation in 30 IMC patients split at the median target-cell fraction among all analyzed cells (15 per group). Median survival is 554 versus 158 days in the higher and lower groups (log-rank *P* = 0.000157). The grouping uses the observed IMC state. **(C)** Workflow. RAMP aligns 165,686 cells from 38 patients using 15 shared proteins. A class-balanced logistic regression learns the spatial target from 5,649 source CD8 T cells, including 478 targets, and is applied to 5,768 CITE-seq CD8 T cells after patient-held-out evaluation. **(D)** Joint UMAP colored by dataset and cell type, using a fixed display sample of 20,000 cells per platform. The ellipse is a visual guide. **(E)** Observed targets and patient-held-out predictions in a selected IMC field. Predictions use the highest-scoring 20% of candidate CD8 cells in the whole patient image. Cyan outlines mark candidates, yellow circles targets or matched predictions, and blue circles unmatched predictions. The field contains 46 candidates, 23 targets and 18 predictions, including 17 true positives. Scale bar, 50 µm; selection details are in Methods. **(F)** AUROC and target enrichment among the highest-scoring 20% for 14 evaluable patients. Median values are 0.707 and 1.98-fold. Standardization and classifier fitting exclude the evaluated patient; RAMP alignment remains fixed. Dashed references are 0.5 and one-fold. **(G)** CITE-seq CD8-only UMAP colored by provider subcluster and transferred score on identical coordinates. Memory/effector cells score above naive cells in all seven patients with both subsets. **(H)** Proteins excluded from alignment and classifier inputs. High is the top score quartile and Lower the remainder within each patient. Points are mean expression ranks for eight patients; lines connect paired values. All four proteins are higher in High in all eight patients. Boxes use the same conventions as the overview (Fig. 1). The CITE-seq scores indicate molecular similarity to the source state and do not assign tissue locations or predict survival in the target cohort.

RAMP aligned 112,497 IMC cells and 53,189 CITE-seq cells from eight additional patients^53^ using 15 shared proteins (Fig. 5C,D). A class-balanced logistic regression learned the spatial target from aligned protein profiles of 5,649 IMC CD8 T cells, including 478 target cells. Standardization and classifier fitting were repeated with each evaluated patient withheld, while the joint RAMP alignment remained fixed. Across 14 patients with sufficient cells and both classes, median AUROC was 0.707 and median target enrichment in the highest-scoring 20% was 1.98-fold; all 14 showed enrichment above one-fold (Fig. 5E,F).

The refitted classifier was then applied to 5,768 CITE-seq CD8 T cells. Memory and effector subclusters scored above naive cells in all seven patients with both subsets, with mean patient-level scores of 0.56 and 0.26, respectively (Fig. 5G). Four proteins absent from the alignment and classifier inputs—CD11a, CD18, CD2 and PD-1—were higher in the highest score quartile in all eight patients (Fig. 5H). Their mean expression-rank module correlated with the transferred score at mean within-patient Spearman *ρ* = 0.480.

CD11a and CD18 form LFA-1, linking the transferred state to an adhesion- and activation-associated program involving CD2 and PD-1. RAMP-enabled transfer thus connected a survival-associated spatial state defined in IMC to a broader protein program in independently profiled, dissociated CD8 T cells.

### Predicted vCAF HLA-ABC state is associated with lung cancer outcome

To connect an unmeasured marker with an existing clinical cohort, we integrated lung CyTOF^49^, mIF^54^ and IMC^50^. A balanced reference contained 1,000 cells from each of 28 stage-matched patients per platform. RAMP aligned these 84,000 cells using nine shared proteins, excluding HLA-A and HLA-ABC (Fig. 6A,B). Measured CyTOF pan-HLA-ABC was converted to a within-patient relative score and transferred to all target cell types by distance-weighted nearest-neighbor averaging. In IMC, we subsequently examined the original authors’ vascular cancer-associated fibroblasts (vCAF; CD146-positive, CD34-negative), retaining their full-panel annotation (Fig. 6C).

**Figure 6:**
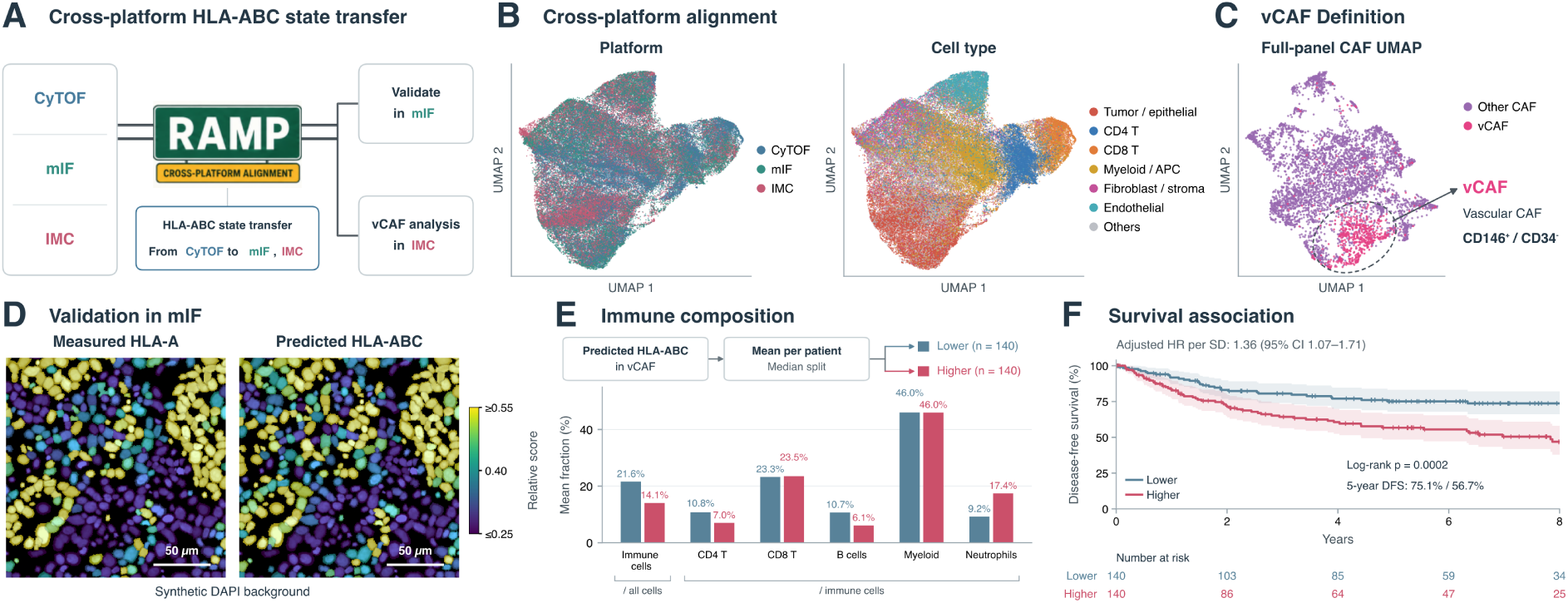
Predicted vCAF HLA-ABC state is associated with immune composition and disease-free survival in lung cancer. **(A)** RAMP integrates CyTOF ^49^, mIF ^54^ and IMC ^50^ using nine corresponding marker features, including the provider-supplied IMC marker combinations specified in Methods. Within-patient CyTOF HLA-ABC relative scores are transferred by weighted nearest-neighbor averaging to all target cell types. HLA-A and HLA-ABC are excluded from alignment. IMC analyses use the original vCAF annotation. **(B)** Joint UMAP of 84,000 cells: 28 stage-matched patients per platform, 1,000 cells per patient, without cell-type quotas. CyTOF reference cells were sampled from a project-curated, QC-filtered reference resource (Methods); IMC and mIF reference cells were retained. **(C)** Separate full-panel UMAP of 5,206 reference IMC CAFs, including 646 vCAF (CD146-positive, CD34-negative) and 4,560 other CAFs. The ellipse is a visual guide; it does not define vCAF membership. **(D)** Measured HLA-A and predicted HLA-ABC relative score in a selected mIF field (389 cells; Spearman *ρ* = 0.870). These are cell-value visualizations using original coordinates, geometry-constrained synthetic cell shapes and a synthetic DAPI background, not acquired fluorescence images. Colors share selected display limits of 0.25–0.55; correlations use unclipped values. Scale bars, 50 µm. Across 693,693 cells from 119 cores and 28 patients, median within-patient correlation is 0.696. **(E)** Immune composition after frozen projection to 2,849,385 IMC cells from 527 patients. The analysis includes 280 eligible stage I–II patients split at the median mean vCAF score (140 per group). Bars are equal-patient mean fractions: total immune cells as a fraction of all cells, and individual immune types as fractions of immune cells. Higher-score patients have lower total immune, CD4 T and B-cell fractions and a higher relative neutrophil fraction. Adjusted tests and all 12 tested endpoints are described in Methods. **(F)** Disease-free survival in the same 280 patients (96 events). Five-year estimates are 75.1% and 56.7% for Lower and Higher (log-rank *P* = 1.92 × 10^−4^); shading indicates 95% confidence intervals. The adjusted continuous-score hazard ratio is 1.36 per standard deviation (95% CI 1.07–1.71, *P* = 0.0104), controlling for age, sex, stage, overall CAF fraction, the global CAF HLA-ABC median score and vCAF fraction among CAFs (Methods).

The mIF cohort provided an auxiliary validation readout because it measured HLA-A, one component of pan-HLA-ABC. Across 693,693 cells in 119 cores from 28 patients, the median within-patient correlation between predicted score and measured HLA-A was 0.696. All core-level correlations were positive (0.275–0.889). In the illustrated core, correlation was 0.827 across 10,719 cells and 0.870 within the displayed 389-cell region (Fig. 6D). HLA-A and pan-HLA-ABC are related but distinct readouts; this comparison supports correspondence of the transferred relative state across platforms.

We extended the frozen alignment and predictor to all 2,849,385 cells from 527 IMC patients, retaining patient-specific maps for the 28 reference patients and fixed platform maps for the remaining patients (Methods). Predictions were averaged over each patient’s annotated vCAF. Among 280 eligible stage I–II patients, the higher-score half had a lower total immune-cell fraction than the lower-score half (14.1% versus 21.6%). Within immune cells, CD4 T and B-cell fractions were lower (7.0% versus 10.8%, and 6.1% versus 10.7%), whereas the neutrophil fraction was higher (17.4% versus 9.2%; Fig. 6E). These differences remained significant after covariate adjustment and correction across the 12 tested composition endpoints.

Higher mean predicted vCAF HLA-ABC was also associated with worse disease-free survival: five-year estimates were 56.7% versus 75.1% in the higher- and lower-score groups (log-rank *P* = 1.92 × 10^−4^). The continuous score had an adjusted hazard ratio of 1.36 per standard deviation (95% CI 1.07–1.71, *P* = 0.0104), controlling for demographic, stage and CAF-related covariates (Fig. 6F). Cross-platform transfer therefore enabled investigation of an unmeasured stromal protein state in an existing clinical cohort, linking molecular information from CyTOF to tissue composition and long-term outcome in IMC.

### Spatial niche transfer identifies a CD39/TIGIT-associated CD8 phenotype

Finally, we used a spatial measurement to screen proteins available only in a larger dissociated-cell panel. In HNSCC CODEX^55^, we quantified CAF/macrophage enrichment among the 15 nearest neighbors of each tumor CD8 T cell relative to its region. A higher patient-average niche score was associated with shorter overall survival (hazard ratio 1.34 per standard deviation, 95% CI 1.03–1.74, *P* = 0.029; 74 patients). This reanalysis builds on the original study’s fibroblast–myeloid niche observations (Fig. 7A,B).

**Figure 7:**
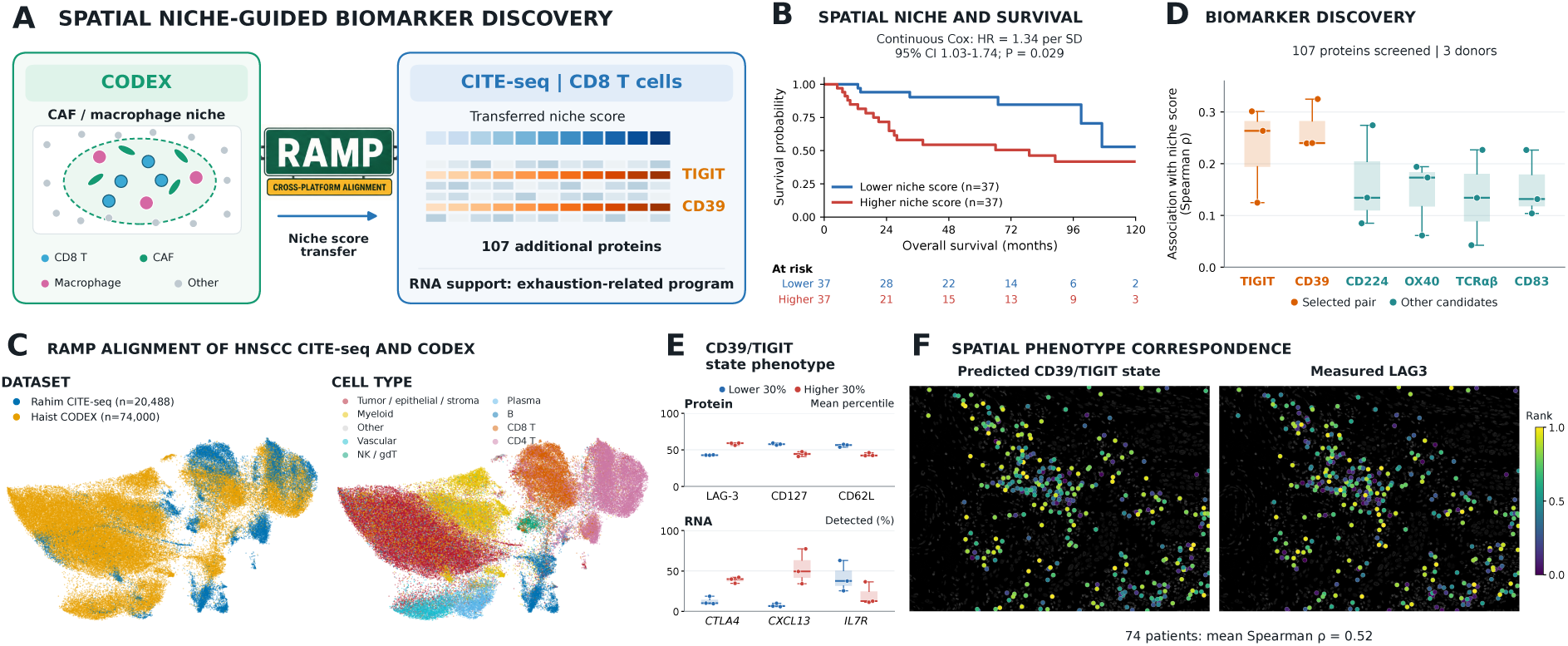
Spatial niche-guided discovery of a CD39/TIGIT-associated CD8 T-cell phenotype in HNSCC. **(A)** A CODEX CAF/macrophage niche score is transferred through aligned proteins to CITE-seq, enabling screening of additional proteins and subsequent molecular characterization. **(B)** Reanalysis of CODEX and clinical data from Haist et al. ^55^. Each CD8 T-cell niche score is the CAF/macrophage fraction among 15 nearest neighbors minus its region-wide background. Kaplan–Meier curves show 74 patients (24 deaths), split at the median patient-average score. The displayed hazard ratio (1.34, 95% CI 1.03–1.74; *P* = 0.029) comes from a univariable Cox model of the standardized continuous score. The niche motivation follows the original paper; these score and survival estimates are from the present reanalysis. **(C)** Joint UMAP of 74,000 CODEX cells (1,000 per patient) and all 20,488 CITE-seq cells from three donors ^56^, aligned on 22 shared proteins. The CODEX maps were extended to all 38,299 primary-tumor CD8 T cells for subsequent transfer. **(D)** Association of six displayed protein candidates with the transferred niche score in 4,788 CITE-seq tumor CD8 T cells. Each point is a donor’s Spearman correlation. CD39 and TIGIT (orange) passed robustness, redundancy and top-5% selection criteria among 107 screened proteins. Neither was measured in CODEX; all six displayed associations are positive in all three donors. **(E)** Characterization using measured CD39/TIGIT state. The bottom and top 30% within each donor contain 1,439 Lower and 1,437 Higher cells; 1,912 intermediate cells are omitted. Points show donor-level mean protein expression percentiles (upper) or fractions with detectable RNA (lower). Higher-state cells have more LAG-3, CTLA4 and CXCL13 and less CD127, CD62L and IL7R in all three donors. Boxes summarize donor values, not confidence intervals. **(F)** The measured CD39/TIGIT state transferred back to CODEX compared with measured LAG3, excluded from alignment and state construction. The selected ROI contains 343 CD8 cells over acquired DAPI; colors are within-patient ranks at cell centroids, not raw marker fluorescence. Across all 74 patients, mean within-patient Spearman correlation is 0.522 (95% bootstrap CI 0.496–0.550). This auxiliary readout was chosen during exploration. The survival association in B pertains to the niche score, rather than the transferred molecular state.

RAMP aligned 94,488 CODEX and CITE-seq cells using 22 shared proteins (Fig. 7C). Patient-specific nearest-neighbor models transferred the CODEX niche score to 4,788 CITE-seq tumor CD8 T cells from three donors^56^. We screened 107 proteins outside the alignment panel for association with the transferred score. CD39 and TIGIT passed the combined cross-donor robustness, redundancy and top-rank criteria (Fig. 7D). Neither was measured in the CODEX panel. CD39 has been associated with tumor-reactive CD8 populations, while TIGIT regulates inhibitory signaling in tumor T cells^57,58^, providing a biological rationale for characterizing their joint state.

We defined this molecular state using the measured within-donor CD39 and TIGIT values, then compared the top and bottom 30% of cells. Higher-state cells had more LAG-3 protein and more frequent CTLA4 and CXCL13 RNA detection, together with less CD127 and CD62L protein and less IL7R RNA (Fig. 7E). All six directions agreed across the three donors. CD127 and IL7R provide concordant protein and transcript readouts of the same receptor. Across the three CITE-seq donors, the selected CD39/TIGIT phenotype therefore showed a coordinated protein and RNA pattern, combining stronger checkpoint-associated features with reduced IL-7-receptor and lymphoid-homing features.

Transferring the measured CD39/TIGIT state back to CODEX yielded positive correspondence with measured LAG3, a marker excluded from both the alignment and state definition. Across 74 patients, mean within-patient Spearman correlation was 0.522 (95% patient-bootstrap CI 0.496–0.550), with spatial correspondence illustrated on acquired DAPI (Fig. 7F). Together, spatial niche transfer identified CD39 and TIGIT within a larger protein panel, connected their joint state to a consistent protein–RNA phenotype, and recovered correspondence with measured LAG3 in the spatial cohort. This workflow demonstrates how spatial context can guide molecular discovery across datasets with complementary marker coverage.

## Conclusion and Discussion

RAMP enables cross-platform integration directly on the protein measurements used for downstream analysis. It returns an aligned cell-by-marker matrix that retains the original marker identities and within-sample expression ordering, allowing marker thresholds and predictive models to operate across technologies. Sample-specific anchoring, cross-platform correspondence and bounded monotone correction jointly support this capability. Across seven integration tasks, RAMP achieved the best overall rank under the main category-balanced summary and under alternative summaries using all evaluated metrics. Fixed-source gates also transferred more accurately after alignment, demonstrating a practical benefit of correcting marker values directly.

Our marker-level analyses show why this output deserves evaluation alongside embedding quality. Correction can improve dataset mixing while changing the relative expression of biologically distinct populations. The examples show reductions or reversals in marker-specific discrimination (Fig. 1), and the broader percentile audit quantifies the magnitude of population shifts. RAMP excludes strict within-sample order reversals by construction. Order preservation protects within-sample marker relationships, while gate transfer tests the cross-platform portability of the resulting scale. Together with population-structure metrics, these evaluations connect the mathematical properties of the transformation to its downstream utility.

Monotone normalization has an established role in proteomics, including ADTnorm’s alignment of antibody-derived tags across CITE-seq studies^39^. RAMP uses cross-platform correspondences to estimate sample- and marker-specific transformations across assays with different measurement chemistries. It complements methods focused on cell matching and shared representations, which can also support imputation and information transfer^31^.

Recent foundation models and pretrained predictors for spatial proteomics support representation learning, marker reconstruction and cell phenotyping. RAMP addresses the complementary need for harmonized marker values that can be used directly in cross-platform analyses. KRONOS learns patch and marker embeddings across fluorescence platforms without an intensity decoder^59^. VirTues reconstructs per-pixel marker intensities across imaging platforms, but standardizes each channel using means and standard deviations computed within each dataset, so the same measured intensity maps to different model-space values in different cohorts^60^. Spatium converts protein abundance to quantile bins computed within each tissue section, so a cutoff on its output denotes a rank within that section^61^. Nimbus generalizes across platforms without retraining but returns marker-positivity probabilities rather than harmonized expression intensities^25^. The fixed-gate experiments demonstrate the practical value of RAMP’s shared normalized marker scale (Fig. 2E,F), while the controlled learning experiments show that aligned measurements can support transferable prediction rules across platforms.

The learning experiments demonstrate benefits across prediction tasks: RAMP led cross-platform annotation in all three model families and achieved the strongest missing-marker results among the tested alignment methods. Percentile preprocessing performed best for the relative-expression prediction target, indicating that the preferred preprocessing strategy can depend on the downstream task.

These findings motivate evaluating RAMP as an input-harmonization or constrained-calibration component for proteomic foundation models. The present evidence establishes benefits for conventional prediction models; effects on large-scale pretraining remain to be tested. Models intended to estimate absolute protein abundance would additionally require experimental calibration targets and independent validation.

The applications demonstrate bidirectional transfer between molecular profiling and spatial cohorts. Reference panels supplied missing marker information to imaging data, while spatially defined states guided molecular characterization in richer single-cell panels. Together, these analyses connected protein programs, tissue context and clinical follow-up across independently collected cohorts.

RAMP currently aligns two or three datasets and depends on informative shared markers, sufficiently overlapping populations and reliable expression measurements. Sparse overlap or unstable low-expression anchors can reduce calibration support. Monotonicity preserves within-sample ordering; preservation of biological differences between samples must also be assessed through cross-platform and downstream evaluations.

The benchmarks use harmonized provider annotations and partially overlapping source cohorts. Metric selection and annotation-class scopes were refined during exploratory analysis, with matched comparisons and full-metric sensitivity summaries documenting the resulting evaluation. The learning experiments assess source-supervised prediction in jointly aligned cohorts; generalization to entirely new target cohorts remains an important next test.

The biological applications establish molecular and clinical associations. Display regions and some score cutoffs were selected during exploration, with cohort-level summaries providing the quantitative evidence. Functional studies are needed to determine whether the checkpoint-associated and stromal states reflect specific mechanisms of immune regulation, and independent measurements and cohorts will be important for validating inferred markers and clinical associations.

Future work can extend RAMP to larger platform collections, quantify uncertainty in cross-platform correspondences and evaluate fixed references in entirely new cohorts. Standardizing the associated prediction workflows would also make it easier to transfer a validated marker or state while retaining its provenance and scale. By combining explicit marker-level alignment with cross-cohort prediction, RAMP connects molecular measurements, tissue organization and clinical follow-up across independently collected studies. This enables existing single-cell and spatial proteomic cohorts to support analyses that require information distributed across technologies.

## Methods

### The RAMP algorithm

#### Input and overview

RAMP aligns protein measurements across two or three datasets while retaining the order of cells along each marker within each sample. The algorithm takes a finite cell-by-marker matrix containing an ordered set of shared proteins, together with dataset and sample identifiers; each sample identifier is unique and nested within its dataset. Cell-type annotations, spatial coordinates and clinical outcomes are not used for fitting. Source measurements are supplied before RAMP-specific preprocessing and may already include processing performed by the original data provider. The implementation described here is RAMP version 2.3.0.

RAMP proceeds in three stages. It first anchors the low-expression population separately for each sample and marker. It then constructs cross-dataset cell correspondences from complementary source-scale and preprocessed representations. These correspondences provide targets for sample- and marker-specific monotone curves; correction strength and the contributions of alternative reference datasets are calibrated on disjoint subsets of the matching cells, and the resulting curves are applied to every cell in each sample.

#### Sample-specific low-expression anchoring

For each sample and marker, negative source values were shifted by adding the absolute minimum plus 0.01; non-negative inputs were left unshifted. An adaptive arcsinh cofactor was defined as the median of the shifted values, or their mean when the median was zero, with a lower bound of 10^−6^. The shifted measurements were transformed as

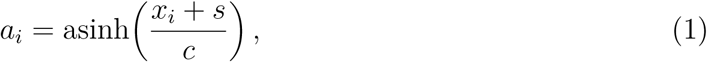

where *s* is the sample–marker shift and *c* the adaptive cofactor. One-dimensional *k*-means with two clusters was fitted to the transformed values using *k*-means++ initialization, 20 starts, at most 500 iterations and a convergence tolerance of 10^−6^. The lower-centroid cluster was accepted as the low-expression anchor when both clusters contained at least max(20, ⌈0.005*n*⌉) cells, the fit converged before the iteration limit, and the centroids were finite and separated by more than 10^−10^. Otherwise the mean of all transformed cells was used as the anchor. The baseline value was

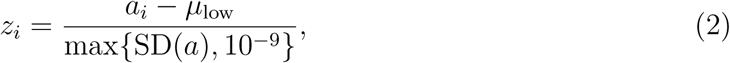

using the population standard deviation within the sample. This places a supported low-expression population at zero and rescales variation without changing within-sample marker order. The low-expression cluster is an algorithmic anchor and does not require a predefined positive/negative gate.

#### Cross-dataset cell correspondences

To balance contributions to matching, the same number of fitting cells was drawn from each dataset. For dataset sizes *N_d_* and sample counts *U_d_*,

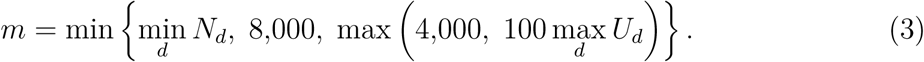

Cells were sampled without replacement using approximately equal sample-level quotas, with unused capacity redistributed among the remaining cells. This internal matching subset is distinct from the population that receives the final transformation.

Two representations were built for each fitting cell, one from its source-scale marker vector and one from its preprocessed vector. Within each representation, marker values were centered across features within the cell and normalized to unit Euclidean length. The resulting vectors were smoothed within each dataset by averaging the cell’s own vector with the mean of its 15 nearest non-self neighbors at equal weight, then normalized again. The two smoothed representations were concatenated and normalized, giving a joint signature for cross-dataset similarity.

Pairwise signature dot products yielded maximum-weight one-to-one assignments via the Hungarian algorithm. For three datasets, assignment weights were attenuated by cycle consistency: cells were followed through the third dataset and back, and the return displacement in each direction was scaled by the corresponding local 15-neighbor radius, giving an attenuation of 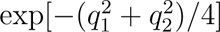 for normalized displacements *q*_1_*, q*_2_. For two datasets the cycle factor is one.

Hard assignments were supplemented by a soft correspondence matrix. Row- and column-normalized softmax probabilities were combined by their geometric mean, with the temperature calibrated so that the median entropy-based effective number of neighbors was 15, using up to 256 evenly spaced rows and columns. Edges in either the top 15 entries of a row or the top 15 of a column were retained. The hard and soft matrices were summed and propagated through Gaussian-weighted within-dataset 15-neighbor transition matrices.

To incorporate marker-order information, each marker was also converted to a within-sample normal score,

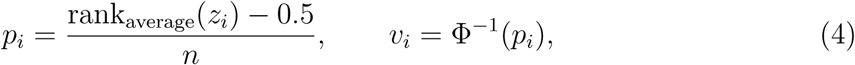

smoothed over 15 non-self neighbors. Propagated correspondence weights were combined with signature similarity and Gaussian support in the smoothed normal-score space by their geometric mean; the Gaussian scale was the median squared normal-score distance over the retained hard and soft edges, bounded below by 10^−8^. Row-normalized correspondence weights then defined barycentric targets as weighted averages of the reference cells’ preprocessed values. Both directions of every dataset pair were retained. Correspondence confidence was 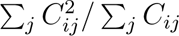 for weighted coupling matrix *C*.

#### Monotone curve fitting and correction-strength calibration

For each directed dataset pair, a deterministic hash of the run identifier, dataset identifiers and cell index allocated fitting cells to curve fitting (12/25), strength calibration (4/25), reference-weight calibration (4/25) or a reserved subset (5/25); the reserved subset entered none of the three calibration stages. Correspondence construction used the complete internal matching subset, so these allocations separate downstream fitting roles within a transductive alignment.

For each query sample, marker and reference dataset, barycentric target values were regressed on the within-sample percentiles *p_i_*. Confidence values were rescaled between their 5th and 95th percentiles, clipped to [0.05, 1] and squared to give fitting weights. Cells were sorted by percentile and divided into at most 20 approximately equal-count bins, requiring at least 15 cells per bin; each bin was summarized by its weighted mean percentile, weighted median target and total weight. Weighted isotonic regression was fitted to these bin summaries and linearly interpolated onto 257 equally spaced percentile positions from zero to one, with tied bin centers combined. A profile required at least four bins and four distinct bin centers; unsupported sample–marker–reference profiles were omitted.

For a fitted reference profile *Q_r_*, the proposed value used during calibration was

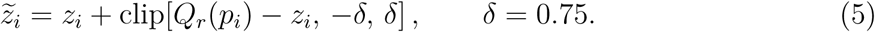

Correction strength was estimated on the strength-calibration subset by fitting the displacement *d_i_* = *z̃_i_* − *z_i_* to the barycentric target *t_i_*:

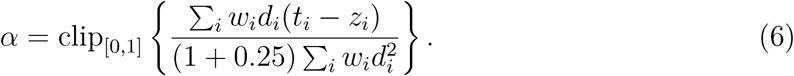

A negligible displacement denominator produced zero strength, and strength was further attenuated when the relative weighted squared-error reduction fell below 0.002. Estimates were computed at both query-dataset and sample level: a sample with at least six calibration cells received weight *n/*(*n* + 30) on its local estimate and the remainder on the dataset estimate, while smaller samples used the dataset estimate alone. This reduces dependence on sparsely supported local fits.

#### Reference weighting and final transformation

Reference contributions were evaluated on the separate reference-weight calibration subset using the reduction in squared error to the barycentric targets. Dataset-level evidence was computed from sample-level weighted mean error reductions, treating samples rather than cells as the aggregation units. Standardized sample-level evidence was shrunk toward the dataset estimate by the same *n/*(*n* + 30) rule, bounded to [−6, 6] before hierarchical combination, and half the combined value was used as the reference log evidence.

A compatibility score measured overlap between each dataset pair. Equal-sized, sample-balanced subsets of up to 3,500 cells per dataset were drawn from the baseline representation; non-constant features were standardized on the concatenated balanced subsets, and PCA to 20 components was applied only when more than 20 features remained. Compatibility combined four coverage fractions by their geometric mean: the fraction of cells with a reciprocal cross-dataset 10-nearest-neighbor relationship in each direction, and the fraction in each dataset whose nearest cross-dataset distance was at most 1.5 times its within-dataset 10-neighbor radius.

For each query sample and marker, reference probabilities *π_r_* were proportional to the exponential of reference log evidence plus log compatibility, and the total alignment weight was 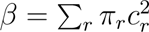 for compatibility scores *c_r_*. Each reference profile was first blended toward the sample’s empirical identity profile using its calibrated strength and the displacement bound; the candidate profiles were then combined with weights *βπ_r_*, with the identity profile receiving weight 1 − *β*. The combined curve was projected onto a non-decreasing function by isotonic regression. If no supported reference profile was available, the baseline values were retained. The final profile *Q* was applied to every cell in the sample using the same displacement-limited operator as during calibration:

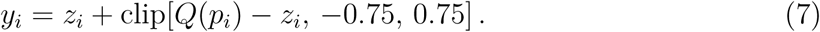

#### Order preservation

Corrections are bounded by 0.75 units on the preprocessed scale. Both the percentile coordinate *p_i_* and the fitted profile *Q* are non-decreasing functions of the input, and equation (7) can equivalently be written as the median of *z_i_* − 0.75, *Q*(*p_i_*) and *z_i_* + 0.75, each of which is non-decreasing in *z_i_*. Consequently, for two cells within the same sample and marker, *x_i_ < x_j_*implies *y_i_* ≤ *y_j_*: the transformation cannot reverse their order, although a non-strict monotone curve may create ties. Order preservation is therefore a property of the map rather than an empirical outcome on a particular cohort. The implementation verifies the displacement bound and the absence of strict within-sample order inversions before returning the aligned matrix.

#### Reproducibility

The algorithm used a base random seed of 42, with deterministic derived seeds for sample selection, *k*-means initialization and fitting-role allocation. The run identifier, cell order, marker order and dataset order were recorded because they determine reproducible fitting behavior. Outputs comprise the preprocessed and aligned matrices together with correspondence, curve-support, correction-strength, evidence and reference-weight records. RAMP returns marker-valued measurements on its preprocessed scale; no separate embedding is required to apply the correction.

### Benchmark cohorts

Four frozen three-platform benchmarks were used, comprising 270,732 cells in 104 dataset-specific sample units. *Bone marrow*: 88,031 cells, 17 units, 14 shared markers with CD19 excluded from alignment — 30,000 Bandyopadhyay CODEX cells (2,500 per donor from 12 donors), 30,000 Levine CyTOF cells (15,000 from each of healthy donors H1 and H2, drawn from the provider’s pre-gated seven-cell-type pool) and all 28,031 Triana AbSeq cells from young healthy samples BM1–BM3; sampling seed 20260820. *TNBC* : 62,701 cells, 11 units, 11 shared markers — 20,000 Tietscher IMC cells from one donor, 22,701 Meyer IMC cells from five coverage-selected patients and 20,000 Shiao CODEX cells (4,000 from each of five donors). *Melanoma*: 60,000 cells, 46 units, 12 shared markers — Hickey CODEX, Hoch IMC and Ptacek MIBI each contributing 20,000 cells from six pretreatment patients, 17 treatment-naive stage III patients and 23 stage III units respectively, balancing patient contributions subject to available cell counts; sampling seed 20260827. The Hickey SOX10/S100 channel serves as the SOX10 proxy. *Lung cancer* : 60,000 cells, 30 units, eight shared markers — Senosain CyTOF, Cords IMC and Monkman CODEX each contributing 20,000 cells as 2,000 from each of ten units, with Senosain patients and eligible Monkman cores selected at random from their frozen parent cohorts and Cords patients by stratified random sampling across six TMA strata; donor seeds 20260828–20260830 and cell seeds 20260838–20260840.

Sampling used no cell-type labels, spatial coordinates or outcomes, except that the Meyer patient selection examined provider labels for B-lineage and endothelial coverage; those labels were not supplied to any alignment method. Platforms are drawn from independent source cohorts, so sample units are not paired patients across platforms and no cross-platform cell-level ground truth exists.

The integration benchmark additionally reuses the frozen lung, melanoma and HNSCC application-cohort alignments as three integration tasks (Fig. 2). The lung story combines Senosain CyTOF, Cords IMC and Zheng mIF; the melanoma story combines Moldoveanu IMC and Engelbrecht CITE-seq protein measurements; HNSCC combines Rahim CITE-seq protein measurements and Haist CODEX. Application sampling is described below. The lung-story task uses the updated Senosain reference described in the HLA-ABC application; all five comparator methods were refitted and all method outputs evaluated with the unchanged protocol. Each comparison method uses the corresponding frozen RAMP cell cohort. These seven tasks can share source datasets and are not seven independent patient cohorts. Complete UMAP and marker-distribution displays are provided for these seven integration tasks (Supplementary Figs. S1–S7); figure-specific display subsets and plotting conventions are given in the supplementary legends below. The 23 primary integration metrics are shown individually (Supplementary Figs. S8 and S9).

Across all seven integration tasks, 303 task-specific sample-unit occurrences correspond to 299 distinct dataset-specific units after accounting for four Senosain units appearing in both lung tasks. This count covers alignment inputs, not all patients in the subsequent full-cohort projections. The seven technologies are CyTOF, IMC, MIBI, CODEX, mIF, CITE-seq and AbSeq; the disease cohorts span breast cancer, melanoma, lung cancer and HNSCC, alongside healthy bone marrow.

### Comparison methods

Comparators used the same frozen cells and shared-marker identities as RAMP. Method-specific input preprocessing was retained rather than imposing RAMP preprocessing on all methods. Consequently, the comparisons evaluate complete configured pipelines. The accompanying preprocessing records distinguish provider processing, additional transformations and correction itself.

#### Harmony and fastMNN

Harmony 1.2.3^29^ was run directly on the source-scale shared-marker matrix without PCA, using dataset as the batch variable, *θ* = 2, *σ* = 0.1, *λ* = 1, up to 100 clusters, at most ten outer iterations and seed 42. A fixed-arcsinh input arm was retained outside the primary six-method ranking. FastMNN used batchelor 1.18.1^30,62^, *k* = 20, cosine normalization, ndist=3, fixed dataset merge order and seed 42. PCA requested 50 dimensions, with the realized dimension limited by marker and cell counts. Integration metrics use the corrected principal components. The separate marker reconstruction used for value-level diagnostics is defined below.

#### cyCombine and CytoNorm

cyCombine 0.2.5^33^ used rank normalization for SOM assignment, an 8 × 8 map, ten training iterations, parametric ComBat, seed 840 and no biological covariate or anchor. This method explicitly supports integration within and across technologies. CytoNorm 2.0.6, with FlowSOM 2.10.0, used the batch-aggregate strategy for data without dedicated technical controls^34,63^. Up to 2,000 cells per sample were pooled into each dataset’s proxy control. FlowSOM used a 15 × 15 map, ten metaclusters, ten training iterations and no additional scaling. Correction used 99 quantiles, their across-batch mean as the goal, monotone splines and seed 42. Official numerical implementations were retained; wrappers handled frozen cell identities and two- or three-dataset inputs.

For cyCombine and fastMNN in the original four benchmarks, inputs followed the saved provider-aware preprocessing: provider CLR for bone-marrow CODEX, arcsinh with cofactor 5 for CyTOF, full-panel CLR for AbSeq, and the distributed analysis scales for already transformed imaging data. TNBC IMC and Shiao CODEX used cofactor-1 arcsinh values. The lung Cords input used the provider’s globally censored, arcsinh-transformed layer, and Monkman CODEX retained its provider transformation and scaling. CytoNorm’s benchmark-only adapter used cofactor-1 arcsinh for bone-marrow CODEX, cofactor-5 arcsinh for CyTOF, log(1 + *x*) for AbSeq, cofactor-1 arcsinh for TNBC and reconstructed Cords counts, cofactor-150 arcsinh for Monkman CODEX, and passthrough of processed melanoma values. These input differences are part of the evaluated pipelines; they were not inferred from the displayed artifacts. The application-task adapters likewise recorded their analysis-ready inputs and source hashes.

#### MaxFuse

MaxFuse^31^ used the same shared proteins as RAMP for its shared feature space and each dataset’s saved full available protein panel for active features. The implementation was pinned to upstream commit 2900b260b697deec678060ea653015291c4dc5cb. Pairwise matching used centroid shrinkage, correlation-based 15-neighbor graphs, three refinement iterations and a 0.3 pivot-filter proportion, with dimensions limited by the available feature ranks. Three-platform integration followed the archived hub-based extension with 15 generalized canonical components. All three hub orientations were evaluated and their metric values combined by the median within each resample; two-platform tasks used pairwise integration. Full active matrices were saved separately from the shared-marker matrices. Integration evaluation used the resulting embeddings rather than an inferred marker-intensity matrix.

### Evaluation

Thirty-one registered metrics were evaluated over 20 deterministic resamples per benchmark, each drawing exactly 5,000 cells without replacement from every source dataset, with samples matched across methods. Seven tasks are evaluated: the four frozen benchmarks together with the lung, melanoma and head-and-neck application cohorts; tasks may share source cohorts. The retained metrics, their grouping into categories and the aggregation rules are defined below. Cell-type metrics use harmonized labels represented in at least two datasets; global mixing metrics retain all sampled cells; silhouette estimates use cell-type-balanced subsamples. Every representation was standardized dimension-wise once over the full benchmark, with no evaluation PCA added.

For the primary summary (Fig. 2), 23 retained metrics are grouped into cell-type structure (5), platform/donor mixing (14), cross-platform label transfer (2) and UMAP (2). This reporting subset and weighting were selected after inspection of the full results and applied identically to every method and task; all 31 original metrics and an equal-individual-metric sensitivity summary are retained. For each task, six methods are ranked per metric according to its direction, with average ranks for ties. Panel D averages those ranks within category, then gives each of four categories 25% weight, and finally averages the seven task scores equally. Category means are not ranked again before aggregation. Panel C instead scales direction-adjusted raw scores to the observed six-method minimum and maximum within each task and metric, then averages metrics within category and tasks equally; an all-equal comparison receives 1 for every method. Panels A and B show Louvain ARI and reference-cell-type UMAP silhouette, respectively, averaging 20 matched resamples from frozen alignments rather than independently refitted models.

#### Metric definitions and clustering

The five cell-type structure metrics were Leiden and Louvain ARI and normalized mutual information (NMI) against harmonized labels, and macro-averaged cell-type silhouette in the corrected representation. Clustering used a symmetrized 15-neighbor graph. For evaluation only, resolution was searched over 10^−5^–8 with 14 geometric bisection steps to approach the number of eligible reference cell types; the partition closest to that count was used. If the count differed by more than max(1, ⌈0.2*K*⌉) for *K* target types, its clustering metrics were marked missing. Labels therefore informed the evaluation cluster count, but not alignment fitting.

The 14 mixing metrics comprised eight cluster-association statistics (ARI and NMI for each of Leiden and Louvain versus dataset and sample unit), dataset and sample-unit neighborhood diversity, dataset neighborhood entropy, a cell-type-conditional mixing acceptance statistic, and unconditional and same-cell-type cross-dataset neighbor fractions. Negative batch-association ARI values were clipped to zero. Neighborhood diversity used the unweighted 30-neighbor inverse Simpson index, normalized as 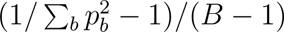 for proportions *p_b_* across *B* batches; this is the registered normalized iLISI implementation, rather than a perplexity-weighted implementation. Entropy was normalized by log *B*. The kBET-like statistic tested 30 same-type neighbors drawn from the first 90 neighbors against the dataset proportions within that cell type, using a chi-squared test with acceptance at *P* ≥ 0.05. Cells with insufficient same-type neighbors were omitted from that statistic. These definitions were identical for all methods.

The two label-transfer metrics were macro F1 and balanced accuracy from 15-neighbor majority voting, holding out each query dataset from the neighbor reference within the frozen aligned representation. The two UMAP metrics were macro-averaged cell-type silhouette and absolute dataset silhouette, with lower absolute dataset silhouette interpreted as better mixing. Silhouettes used balanced samples across eligible groups, normally capped at 3,000 cells. The complete 31-metric registry and task-level scores are supplied as supplementary CSV tables. The accompanying `rank_sensitivity.csv` compares equal-category and equal-metric summaries for both the main subset and the complete registry; RAMP remains first in all four summaries.

#### Fixed-source marker thresholds

Fixed-source marker-threshold evaluation retains the original four-benchmark scope (Fig. 2E,F). Eligible units contain at least ten reference-positive and ten reference-negative cells for a marker task, and source calibration requires at least 50 cells per class. Each method calibrates one source-platform threshold to maximize balanced accuracy, weighting classes equally and units equally within each class; ties favor higher specificity. The threshold is frozen and applied to each other platform, with no target-label threshold selection. CD3 contrasts T with definite non-T cells; CD4 contrasts CD4 T with CD8 T; CD8 contrasts CD8 T with annotated CD4/CD8-negative T cells. Ambiguous and non-joint labels are excluded. Target-unit BA is averaged within direction, then across six directions per benchmark–marker task. For the overall score, tasks are averaged within benchmark and four benchmarks are weighted equally. All 11 eligible tasks are shown. Raw is an independent baseline. Standard PCA-fastMNN reconstructed marker values use the same gating protocol; its corrected PCs remain in A–D. MaxFuse has no marker output. This is threshold transfer on jointly aligned frozen data, not a held-out alignment-model evaluation. Illustrative densities use asinh values (cofactor 1) and Gaussian KDE with bandwidth factor 0.22, balancing classes and donors; source and target share plotting ranges within method and marker. No task-level significance tests are reported.

### Cross-platform learning experiments

#### Data, information boundaries and preprocessing

For the cross-platform learning experiments (Fig. 3), cells, markers and input transformations were recorded in versioned task manifests. The updated Lung reference in B,C contains 84,000 cells: 28,000 each from Senosain CyTOF, Cords IMC and Zheng multiplexed immunofluorescence, with 1,000 cells per registered patient. The Senosain cells were resampled from the reviewed 140,000-cell annotation pool; Cords and Zheng were unchanged. Patient identities and sample units were registered separately; a tissue region was not treated as an independent patient. All methods used the same unlabeled input pool within each experiment. Classification scope restrictions affected supervised training, validation and target scoring, but did not remove cells from the alignment pool. Target answers did not enter alignment or prediction-model fitting and selection. Predictions were saved with cell identities and checksums before target scoring.

The updated annotation comparison uses the following task-specific scopes: bone marrow, CD4 T, CD8 T, B and monocyte; TNBC, CD4 T, CD8 T, myeloid, stroma and tumor/epithelial; original three-platform melanoma, myeloid, stroma and tumor; original lung benchmark, CD8 T, endothelial, myeloid and tumor/epithelial; Lung story, CD4 T, CD8 T, myeloid/APC and tumor/epithelial; Melanoma story, B lineage, CD4 T including Treg, and CD8 T; and HNSCC, B, CD4 T and CD8 T. Cells outside these sets were excluded from supervised classification rather than pooled into an “other” class. The scopes were revised after exploratory review of the initial results, annotation compatibility, marker support and training coverage; they were not prespecified before the initial benchmark. All methods were evaluated with the same revised scopes and matched source draws. The bone-marrow experiment was unchanged. Original full-scope scores are retained with the source data. Shared classes omitted for insufficient training coverage, such as CD8 T in the original three-platform melanoma benchmark, are not claimed to be biologically absent or intrinsically incomparable. Missing-marker prediction uses all frozen cells without the annotation-class restrictions.

We compared ten pipelines: unaligned assay-ready input, RAMP 2.3.0, Harmony, cyCombine, CytoNorm, fastMNN, MaxFuse, upper-clipped *z*-score, min–max and within-unit percentile transformation. The existing numerical alignment implementations were retained, with dataset-specific input contracts distinguishing source-scale input for RAMP and Harmony from assay-ready input for other methods. Assay-ready transformations included the registered arcsinh, log(1 + *x*) or shared-panel CLR operations and passthrough of provider-processed values where appropriate. This comparison evaluates the configured pipelines, including their preprocessing. Clipped *z*-scores cap each marker at its within-unit 99th percentile and then standardize it within each dataset. Min–max scaling maps each dataset– marker range to [0, 10]. Percentile transformation uses (*r* − 0.5)*/n* within each unit, where *r* is the average rank for ties and *n* is the number of cells. Constant columns are set to zero for the first two strategies and to 0.5 for percentile transformation.

FastMNN and MaxFuse contribute their corrected embeddings to the same downstream prediction models; they are not required to return marker values. Unlike the full-active-panel MaxFuse configuration used for integration benchmarking (Fig. 2), this experiment uses the same shared-marker panel for its shared and active feature spaces. Features are standardized within dataset, and the first registered dataset is the fixed hub for three-view integration, with dimensions limited by the retained marker rank. RAMP uses its fixed parameters and seed 42. Method configurations, input contracts and the frozen experiment profile accompany the source data.

#### Source supervision and prediction models

The primary single-source annotation and missing-marker comparisons use 500 source training cells and 512 source validation cells. Approximately 20% of source patients are reserved for validation where multiple verified patients are available; single-patient sources use disjoint cell sets. Where patient crosswalks are unresolved, splits and scores are explicitly sample-unit descriptive. Cell budgets are drawn as prefixes of seeded random permutations, independently of cell-type labels for these two tasks. The same cells and candidate grids are used across processing methods. Three seeds, 20260919, 20260920 and 20260921, govern source sampling and prediction-model initialization. Infeasible budgets or splits are registered before model comparison and excluded identically across methods, without inspecting target performance. These budget and split rules are distinct from the exploratory class-scope revisions described above.

Linear models are logistic regression for annotation and ridge regression for marker prediction. Their inverse regularization strength *C* or ridge penalty *α* is selected from {0.1, 1, 10}. Nearest-neighbor models use *k* ∈ {5, 15, 50} and uniform or inverse-distance weights; *k* is capped by the training-cell count. Neural networks have one ReLU hidden layer, width 32 in the primary comparison, Adam learning rate in {0.001, 0.01} and *L*_2_ penalty in {0.001, 1}. Training uses batches of at most 128 cells, at most 100 epochs and early stopping after 15 epochs without improvement on source validation. The best source-validation checkpoint is retained. Each model family also compares unchanged features with a global feature-wise standardization fitted only on source training cells and applied unchanged to source validation and target cells. Hyperparameters are selected by source-validation macro-F1 for classification and within-unit Spearman correlation for regression; ties follow a fixed lexicographic ordering of the effective configurations. No target answers select models or scaling.

#### Marker masking and relative-expression targets

One marker is held out at a time from every platform, including aliases of the same physical marker. Input operations that depend on the marker panel are recomputed after masking, and all representations, including RAMP matching and transformation, are refitted using the remaining markers. Thus, full-panel alignment caches do not supply information about the withheld marker. The source retains its measured marker values only as supervised answers. Before training/validation splitting, 64 additional cells per source unit are reserved for calibration and kept separate from both sets. Their empirical distribution converts source marker measurements to relative-expression targets using a mid-distribution empirical cumulative distribution function. Calibration is unit-specific and is not fitted from target measurements. Calibration cells can come from source validation patients, but their cell IDs are disjoint from the model-training and validation sets. Marker holdout begins at the frozen provider-processed matrix: upstream denoising, segmentation, QC and sampling were not rerun with the marker absent. The Senosain expression layer is provider-normalized/denoised, rather than raw instrument output. Marker holdout for the Lung configuration uses these mapped channel labels: CD31, EPI and CD45 include the provider-supplied Cords marker combinations specified in the lung application Methods. The regression target is the relative expression of the corresponding measured channel in each platform.

For scoring only, target truth is defined as (*r_i_* − 0.5)*/n_u_* from the measured marker within target unit *u*. MAE is computed against this common target after clipping predictions to [0, 1]. Spearman correlation uses the un-clipped predictions and measured values within units containing at least 30 cells and nonconstant truth. Constant predictions receive effective correlation zero; units with insufficient or constant truth support do not contribute to the correlation but remain eligible for MAE. Units are averaged within each patient and patients equally, with sample-unit summaries retained where patient identity is unresolved. This task evaluates relative expression and ordering, not recovery of absolute protein concentration. It covers 79 eligible benchmark–marker combinations and 28 directed transfers across six tasks. The original three-platform melanoma benchmark is excluded from strict marker holdout because upstream marker dependencies in its distributed MIBI input could not be resolved. This eligibility restriction applies to every method. A source split with no varying marker truth for validation is likewise excluded uniformly.

#### Scoring, aggregation and source records

Classification uses a confusion matrix normalized by each target patient’s cell count and then averaged equally across patients, or across registered sample units for unit-descriptive data. Macro-F1 and balanced accuracy are computed from this matrix across the fixed task ontology; undefined class ratios are assigned zero. Accuracy is retained in the accompanying tables. Within a model family, results are averaged over available registered seeds, then over held-out markers within direction for regression, then directions within benchmark, and finally benchmarks equally. B,C additionally average the three model families equally for each benchmark point. Boxplots are descriptive summaries over overlapping benchmark configurations; random seeds and configurations are not independent-patient replicates. Per-model scores, source/target mappings, eligibility records and input provenance are supplied under supplementary_data/trainability/. Model-specific annotation results are shown for every benchmark (Supplementary Figs. S10 and S11); missing-marker Spearman and MAE are shown separately (Supplementary Figs. S12 and S13).

### Within-sample marker-order audit

In an earlier audit of marker-valued output, 1,151 sample × marker combinations across the four benchmarks were tested. A combination counts as affected when at least one cell pair satisfies input *x_i_ < x_j_* and output *y_i_ > y_j_* with a reversal margin exceeding 10^−12^; pairs tied on input are excluded. The audit records the existence of a reversal per combination, not the number of reversed pairs or the severity of any resulting biological artifact. The original audit excluded fastMNN and MaxFuse because their integration embeddings do not have measured-marker axes; the additional fastMNN reconstruction diagnostic below is separate.

For the sample-level specificity examples, marker values were converted separately for each method to percentile ranks among all cells of the named sample as average rank*/*(*n* + 1), and AUROC was computed between the two named populations within that sample. The CD4 example uses CyTOF H1 (4,029 CD4 T, 2,883 CD8 T, 2,756 B and 3,608 monocyte cells); the CD8 example uses Shiao CODEX h17 (164 CD4 T, 120 CD8 T and 420 macrophage cells); and CD14 uses CyTOF H2 (4,264 CD4 T, 3,563 CD8 T, 2,185 B and 2,600 monocyte cells).

#### Cell-type marker distribution shifts

For the cell-type marker-shift audit (Fig. 1D), the common baseline was the frozen raw marker matrix. Raw and corrected values were ranked independently among all cells in each sample and marker, with average ranks for tied values, and converted to 100 × rank*/*(*n* + 1). For each sample × cell type × marker combination, we computed the signed and absolute changes in the 25th percentile, median and 75th percentile, using linear quantile interpolation. The displayed statistic was the absolute median change in percentile points. The same 6,173 combinations from 103 eligible units out of 104 loaded sample units were used for RAMP, Harmony, cyCombine, CytoNorm and reconstructed fastMNN, retaining all shared markers and resolved groups with at least 50 cells per sample. Smaller groups and unresolved/non-joint labels were omitted from the summary but retained in the sample-wide ranking reference and complete tables. Combinations were weighted equally; benchmarks, patients and cell counts were not separately reweighted, and no significance tests treated these combinations as independent patients. Changes relative to method-specific preprocessed inputs were retained separately. The display range was 0–60 points; observations above 60 were represented by right-pointing boundary markers, while box and whisker statistics used full values. Changes in ties can alter average-rank percentiles without a strict order inversion. Cell-type labels were used only for evaluation, not for alignment.

For the marker-coordinate analyses (Figs. 1C,D and 2F), fastMNN marker coordinates were reconstructed as corrected principal-component scores multiplied by the transpose of the saved rotation matrix, without restoring original-scale centers or inverting cosine normalization. This is the standard PCA reconstruction, not the *d* = NA experiment, and serves as a marker-coordinate diagnostic; integration metrics retain corrected principal components. MaxFuse has no analogous corrected marker output.

### Display settings

#### Ridge plots

All cells enter the histograms (Fig. 1B; Supplementary Figs. S1–S7). For each method–marker, bin width is the pooled 0.5th–99.5th percentile span divided by 160. Gaussian smoothing uses sigma 6 bins, truncation at four standard deviations and constant boundary handling. Grids cover all observed values plus at least four bandwidths at each end. Curves are peak-normalized; height does not represent cell number. Compact axes in Figure 1B and selected supplementary rows (specified in their legends) show the union of displayed sample-curve ranges above 0.5% of peak height, padded by 4% within the grid. RAMP SOX10 has a −1.5 left limit in Figure 1B and Supplementary Fig. S3; other supplementary axes show the full grid. All values contribute to density estimation, including tails outside compact windows. Raw values in Figure 1B use asinh(*x/*1); supplementary transformations are specified in their legends. Each method retains its output scale, so horizontal values are not comparable across method rows.

#### UMAP

Each representation was standardized dimension-wise across the full benchmark, without additional PCA, and embedded with 30 neighbors, minimum distance 0.15, Euclidean distance, spectral initialization and seed 42. Cell-type annotations were attached after fitting, for coloring only. Harmonized labels are project harmonizations of provider annotations rather than a single provider ground truth. UMAP display refinements alter point rendering and layer order only and are applied identically to all methods within a panel.

### Bone-marrow CD19 completion and validation

#### Inputs and channel-failure detection

The frozen 88,031-cell cohort and sampling scheme are described under Benchmark cohorts. Fourteen functioning shared markers were used: CD117, CD11b, CD11c, CD123, CD14, CD3, CD33, CD34, CD38, CD4, CD45, CD45RA, CD61 and CD8. CD19 was retained only as a reference-platform target and as an observed CODEX channel for evaluation. For label-free anomaly detection, reference-platform ExtraTrees models predicted each marker from within-donor ranks of the remaining markers. Prediction–measurement disagreement and distribution diagnostics were converted to within-donor percentiles, and the two largest were averaged. Anomaly-rank counts allowed ties. Cell labels were used afterward to assess observed CD19 discrimination between B and CD4/CD8 T cells.

#### Masked-marker neural network

The functioning marker matrix underwent RAMP 2.3.0 low-expression preprocessing (KMEANSLOW), followed by pooled standardization. A fixed, donor-balanced sample of 4,000 cells per platform supplied 12,000 training cells. Each training example paired a cell with a sampled target marker; when that marker was an input, its value was masked and a binary mask vector was concatenated to the features. The shared trunk had fully connected layers of 64 and 40 units with SiLU activations. Each platform had a 40–8–40 input adapter with a tanh bottleneck; each marker had a 40–16–1 prediction head. Platform-specific 40–8–target output residuals and input adapters were scaled by 0.20. All three platforms supplied functioning-marker targets, but only CyTOF and AbSeq supplied CD19 targets.

Training used AdamW with learning rate 10^−3^, weight decay 2 × 10^−4^, batch size 512, 60 epochs and gradient clipping at norm 5. The objective was smooth L1 loss with transition parameter 0.5, weighted to balance the two unsupervised expression clusters within each donor and target marker. Seeds and the selected cells were fixed in the saved model. CODEX predictions were the equal mean of the CyTOF- and AbSeq-specific output heads. The completed KMEANSLOW-scale matrix then entered RAMP’s matching and correction backend directly, without a second preprocessing transformation.

#### Permanent-marker holdout and summary statistics

Each of the 14 functioning markers was removed permanently from every platform’s input features and from CODEX targets, while remaining available as a target in the two reference platforms. A separate model was trained for each task using the same architecture and training protocol. This is marker holdout on the frozen donor cohort. Spearman and Pearson correlations compared predicted and measured KMEANSLOW values within each donor. For AUROC and average precision, measured values were partitioned by two-cluster *k*-means within donor and marker, with the higher-centroid cluster designated positive. Average precision was non-interpolated. The four displayed donor summaries average the 14 marker-specific values within each donor; medians were then taken over the 12 donors. The CD3 scatter uses all 30,000 CODEX cells and a pooled correlation. CD3 was selected after reviewing the holdout tasks as a familiar example consistent with the accompanying UMAP.

The CD3 UMAP was fitted jointly on all 88,031 cells using the 13 remaining shared markers, with CD3 and CD19 excluded. The predicted-CD3 color scale used the 1st and 99th percentiles of measured CODEX CD3. UMAP comparisons without and with completed CD19 used condition-wise pooled standardization, no PCA, 30 neighbors, minimum distance 0.15, Euclidean distance and seed 42. The B-versus-NK silhouette used the same balanced sample of 2,500 cells per group. CD19 ridge densities used bandwidth factor 0.35 and were independently peak-normalized.

#### Spatial display

The fixed preprocessing and predictor were applied to all 83,384 H14 atlas cells. Panels A and G use the same field and orientation, with acquired DAPI, acquired CD19 in A and the provider’s registered whole-cell masks in G. A 1,024-pixel parent ROI was chosen as the highest-F1 window among 1,331 eligible windows using predicted CD19 *>* 1.4 versus B-cell annotation. The displayed 512 × 512-pixel field was subsequently selected for maximum centroid density within that ROI using a 16-pixel stride. The display therefore illustrates correspondence in a selected field; whole-cohort B-identity AUROC was computed separately within each of the 12 sampled donors and averaged equally. The continuous overlay is neutral gray at or below 1.0 and saturated yellow at or above 1.5 on the prediction scale, with original DAPI brightness providing texture. These are visualization limits, not positivity or non-expression definitions. Scale bars are 50 µm.

### Melanoma spatial-state transfer

#### Cohorts, target and survival

The IMC input included all 30 patients with provider cohort field ICI; the CITE-seq input included all eight StageIII patients. RAMP aligned 112,497 IMC and 53,189 CITE-seq cells using PDL1, CD45, CD4, CD20, CD8, CD45RA,

CD40, CD3, CD45RO, CD14, CD31, CD29, LAG3, ICOS and OX40. The source candidate pool contained 4,588 provider-annotated Tc.ae and 1,061 Tc.naive cells from 28 patients. The target comprised Tc.ae cells with Ki67 positivity probability ≥ 0.5 and nearest melanoma centroid distance ≤ 30 µm. The target definition was fixed before transfer. Survival analysis used all 30 IMC patients, splitting the observed target-cell fraction among all analyzed cells at its median (3.042 × 10^−4^). Overall survival from immune-checkpoint-therapy initiation was compared by a two-sided log-rank test.

#### Classifier and patient-held-out evaluation

A StandardScaler followed by logistic regression used *C* = 0.5, balanced class weights, the liblinear solver, at most 5,000 iterations and seed 20260829. Ki67, spatial distance and survival outcomes were excluded from classifier inputs. Each evaluated patient’s cells were excluded from scaler and classifier fitting; the jointly fitted RAMP representation remained fixed. AUROC and enrichment were summarized for 14 patients with at least 20 candidate cells and both target classes. Enrichment was the target fraction among the highest-scoring 20% divided by the target fraction among all candidate cells in that patient. After evaluation, the model was refitted on the complete source candidate pool and applied to 5,768 provider-defined CD8_T CITE-seq cells.

The illustrative spatial field came from patient 22RD and was selected after ROI and score-cutoff exploration. Predictions were called in the highest scoring 20% of all 280 candidate CD8 cells in the whole patient image before restriction to the displayed field. The field contained 46 candidates, 23 targets and 18 predictions, of which 17 were targets. Cohort-level held-out performance is reported independently of this field selection.

#### Target-cohort protein characterization and display

CD11a, CD18, CD2 and PD-1 entered neither alignment nor classifier inputs. For each cell, log(1 + ADT) was centered by the mean over all 130 non-isotype ADTs. These values were average-ranked within patient and marker and divided by *n* + 1. High-score cells were those with within-patient score rank ≥ 0.75; Lower comprised the remainder. Each plotted point is a patient’s group mean. The four-protein module was the mean of their ranks, and its reported correlation was the equal-weight mean of eight within-patient Spearman correlations.

The joint UMAP display sampled 20,000 cells per platform: 666–667 per IMC patient and 2,500 per CITE-seq patient, with seeds 20260820 and 20260821. Joint and CD8-only UMAPs used standardized aligned proteins, 30 neighbors, minimum distance 0.20 and Euclidean distance, with seeds 20260820 and 20260829. CITE-seq views used the same distance-preserving rotation for layout. Ellipses were visual annotations and did not define model targets.

### Lung vCAF HLA-ABC inference and survival analysis

#### Balanced reference and marker transfer

All 28 eligible Zheng mIF patients were retained. Senosain CyTOF and Cords IMC patients were selected to match their stage distribution (19 stage I and nine stage II per dataset) while spreading across acquisition dates or tissue microarrays. Each patient supplied 1,000 cells without cell-type quotas, giving 84,000 reference cells. For this application, the Senosain reference was drawn from a project-curated 140,000-cell reference resource: the first 1,000 QC-kept cells per selected patient in a frozen random order were used. Mixed annotations passing QC were retained, and neither cell identities, HLA values nor outcomes were used to set sampling quotas. The patient list and the original Cords and Zheng reference cells were unchanged. This reference also supplies the lung-story integration benchmark (Fig. 2) and its updated single-source annotation and missing-marker experiments (Fig. 3B,C). Alignment used nine harmonized feature labels: CD3, CD4, CD8, CD31, Ki67, vimentin, EPI/cytokeratin, CD45 and HLA-DR. These labels map to platform-specific channels. In Cords IMC, the features labeled CD31, EPI and CD45 were read from the provider’s “vWF + CD31”, “Pan Cytokeratin + Keratin Epithelial” and “CD45RA + CD45R0” rows of the counts assay, respectively. The corresponding Senosain channels were CD31, Cytokeratin and CD45, and the Zheng channels were CD31, Pan-Cytokeratin and CD45. The Cords combinations were supplied in the deposited data; we did not construct them by adding separate channels. Thus, alignment used corresponding markers or marker combinations rather than assuming identical antibody targets across platforms. The complete nine-feature mapping is supplied in supplementary_data/lung_marker_channel_mapping.csv. HLA-A and HLA-ABC were excluded from alignment.

Within each source patient, CyTOF HLA-ABC was converted to average rank divided by *n* + 1, where *n* is the number of reference cells. Predictions were normalized weighted means of these scores over the 30 nearest source cells in aligned protein space. Feature standardization used source-cell means and standard deviations. Weights were proportional to exp[−(*d/d*_30_)^2^], where *d*_30_ is the query’s 30th-neighbor distance; *k* = 100 was retained as a sensitivity configuration. Prediction included all cell types before restricting to author-annotated vCAF. Measured mIF HLA-A was converted to a relative score across all available cells per patient; predictions were not re-ranked. Correlations used unclipped cell-level values and were summarized separately by core and by patient.

The joint UMAP used standardized reference features, 30 neighbors, minimum distance 0.12, Euclidean distance and seed 42. The CAF-only UMAP used the original 41-protein Cords panel, standardized over 5,206 reference CAFs; vCAF membership came from the authors’ CD146-positive, CD34-negative annotation. The mIF spatial panel displays 389 cells from a selected region of core C_30, patient P_26. Shared synthetic cell shapes and a synthetic DAPI background were constrained by the original centroids and measured nuclear geometry. These renderings visualize measured and predicted cell values and are not acquired fluorescence images. A selected common range of 0.25–0.55 was used for color saturation only.

#### Projection to the complete IMC cohort

The 28 reference patients retained their patient-specific RAMP profiles. For the remaining 499 patients, the joint alignment was not refitted. Instead, one monotone platform map per marker was estimated by isotonic regression between paired baseline and aligned values from all 28,000 reference IMC cells. Interpolation retained the endpoint displacement outside the observed range, blended 10^−6^ of the identity map, and clipped displacement to ±0.75. Each new patient’s initial KMEANSLOW normalization was fitted on 1,000 fixed randomly sampled cells and then applied to every cell before the platform map and frozen HLA-ABC predictor. Thus, the 1,000-cell samples fitted normalization; predictions covered all 2,849,385 cells from 527 patients. This fixed-map projection is an application-specific extension of RAMP.

#### Patient summaries and statistical analysis

The patient score was the mean prediction across all annotated vCAF, pooled over available cores. Eligibility required provider stage group I–II, at least 15 vCAF, at least 50 CAFs, positive and valid DFS follow-up, a binary event indicator and complete model covariates. No proximity-to-T-cell or SMA-CAF requirement was imposed. This gave 280 patients and 96 events. Higher and Lower groups were defined by the median patient score (0.63407; 140 patients each), without optimizing the survival cutpoint.

Composition was summarized as equal-patient mean fractions, with total immune cells divided by all cells and immune subsets divided by immune cells. Two-sided linear models adjusted for tissue microarray, age, sex and stage, using HC3 standard errors; Benjamini– Hochberg correction covered all 12 composition endpoints. The adjusted *q* values for total immune, CD4 T, B-cell and neutrophil fractions were 2.03 × 10^−4^, 6.32 × 10^−4^, 5.49 × 10^−4^ and 6.32 × 10^−4^, respectively.

DFS in days was converted to years by division by 365.25. Kaplan–Meier curves and five-year survival estimates used the median groups; a two-sided log-rank test compared them. Cox regression used the continuous vCAF score, standardized within the analysis cohort, with Efron handling of ties. The model adjusted for age in years, sex, categorical provider stage, overall CAF fraction, the global CAF HLA-ABC median score from the present three-platform analysis, and vCAF fraction among all CAFs. Continuous score and fraction covariates were standardized. Schoenfeld-residual diagnostics and analyses excluding reference patients were retained with the model outputs. These are exploratory association analyses, not independent clinical validation.

### HNSCC niche-guided screening and spatial phenotype correspondence

#### Alignment and spatial query

CODEX contributed 1,000 primary-tumor cells from each of 74 patients, sampled after pooling their ROIs (seed 20260823). All 20,488 CITE-seq cells from three donors were retained. The 22 shared proteins were CD45, CD3, CD7, CD4, CD8, CD25, CD20, CD38, CD56, CD57, CD16, CD11c, CD11b, HLA-DR, CD21, CD31, CD69, ICOS, PD1, CD45RO, CD45RA and PD-L1. Joint UMAP used standardized aligned values, 30 neighbors, minimum distance 0.25, Euclidean distance and seed 20260824. Cell annotations were added after alignment.

For each CODEX primary-tumor CD8 T cell, the niche score was the fraction of CAFs and macrophages among its 15 nearest non-self neighbors minus their fraction among all cells in the same ROI, excluding the index cell. Patient scores were unweighted means over evaluable CD8 cells. The 74-patient survival analysis included 24 deaths. Median splitting gave 37 patients per curve; the reported hazard ratio and two-sided Wald test came from a univariable Cox model of the standardized continuous score.

#### Score transfer

Empirical monotone maps from source to RAMP values were fitted for each CODEX patient and shared marker and extended to all 38,299 primary-tumor CD8 T cells, using constant boundary extrapolation. Feature scaling combined equal-patient CODEX and equal-donor CITE-seq first and second moments, giving each platform half the total weight. One *k* = 30 nearest-neighbor predictor was fitted per CODEX patient, using all available cells when fewer than 30 were present. For each CITE-seq query, neighbor weights were proportional to exp(−*d/d*), where *d* was the median neighbor distance. The cell-wise median over the 74 patient-specific predictions defined the transferred score in 4,788 CITE-seq tumor CD8 T cells. Survival outcomes were not used for transfer or protein screening.

#### Protein screening and molecular characterization

The 107 candidate proteins excluded the 22 shared markers, seven isotype controls and LAG-3. Four association views were evaluated within each donor: raw ranks; log-counts centered by the within-cell mean over 129 non-isotype ADTs excluding LAG-3; partial Spearman correlation adjusting for mean log-isotype counts; and partial Spearman correlation adjusting for the marker-specific mean of the other 128 background ADTs. Donors were weighted equally. Candidates required positive CLR and isotype-adjusted associations in all three donors, a positive global-ADT-adjusted association in at least two, median rank at most 15 across views, and positive mean associations after leaving out each donor. Redundancy filtering removed candidates with absolute equal-donor correlation ≥ 0.85 to a higher-ranked candidate. The final compact set additionally required a top-5% rank among all 107 proteins.

The measured CD39/TIGIT program was the mean of their within-donor CLR expression percentiles. For group comparisons this mean was re-ranked within donor, and the bottom and top 30% were retained. The 30% fraction was chosen after fixing the six characterization readouts and comparing 10, 20, 25, 30, 40 and 50%; all six contrasts had consistent directions across donors at each fraction. Protein summaries were mean within-donor expression percentiles, with CD39 and TIGIT excluded from the normalization background. RNA summaries were the fraction of cells with a nonzero count. The plots show three donor-level summaries per group and do not treat cells as independent replicate patients.

#### Return transfer and LAG3 correspondence

The fixed measured CD39/TIGIT program was transferred from CITE-seq to CODEX using one distance-weighted *k* = 30 predictor per CITE-seq donor and the cell-wise median of the three predictions. Feature scaling and CODEX maps remained fixed. LAG3 entered neither the shared-marker alignment nor the program definition. Spearman correlations between predicted program and measured LAG3 were computed within all 74 CODEX patients and averaged equally; the 95% interval was obtained by resampling patients. LAG3 was selected as an auxiliary readout during exploration. The displayed patient-29770 ROI was the highest-correlation region among those with at least 200 CD8 cells; it contained 343 CD8 cells and had *ρ* = 0.680. Both overlays use within-patient percentiles at the original cell centroids over acquired DAPI. The footer reports the all-patient mean correlation, independently of ROI selection.

## Data Availability

This study reanalyzes publicly available data; no new experimental data were collected. Derived scores and benchmark summaries were generated computationally. All datasets are available from their original publications, which should be cited by anyone reusing them. The four benchmarks and four application cohorts draw on 17 datasets reported in the following studies and data records: breast cancer imaging mass cytometry^43,44^ and CODEX^45^; healthy bone marrow CODEX^46^, mass cytometry^47^ and AbSeq^48^; lung cancer mass cytometry^49^, multiplex immunofluorescence^54^, imaging mass cytometry^50^ and CODEX^51^; melanoma CODEX^40^, imaging mass cytometry^41,52^, multiplexed ion beam imaging^42^ and CITE-seq^53^; and head-and-neck squamous cell carcinoma CODEX^55^ and CITE-seq^56^. Where those studies deposited data in a public repository, we used the deposited version: bone marrow CODEX images and segmentation masks are available at figshare+ (https://plus.figshare.com/articles/dataset/Normal_Bone_Marrow_CODEX_Images/25122311), the head-and-neck CODEX data at Mendeley Data (https://doi.org/10.17632/t4k5kj3cxr.1), the head-and-neck CITE-seq data at NCBI GEO under accession GSE212797, the melanoma CODEX data at Dryad (https://doi.org/10.5061/dryad.k0p2ngfcc), the melanoma multiplexed ion beam imaging data at Mendeley Data (https://doi.org/10.17632/79y7bht7tf.2), and the lung multiplex immunofluorescence data at Dryad. For the remaining cohorts, access follows the terms stated in the corresponding publication. Alignment and prediction use provider-processed cell-level matrices. Acquired image channels and segmentation masks were additionally used for the specified spatial displays. Input layers and subsequent transformations are described in Methods and the accompanying preprocessing records. The supplementary data include the metric registry, all seven-task metric scores, alternative ranking summaries, per-benchmark and per-model prediction scores, and frozen protocol records underlying the cross-platform learning experiments (Fig. 3). Clinical follow-up data were used as distributed by the original studies, and no attempt was made to re-identify any individual.

## Code Availability

The public repository for the RAMP software and the study analysis scripts is https://github.com/QunzhiXu/RAMP.

## Acknowledgments

This work is supported by the startup funding to Y.Z. by the University of Texas MD Anderson Cancer Center. We thank the investigators of the studies listed under Data Availability for making their data publicly accessible.

## Author Contributions

Y.Z. conceived the project. Q.X., T.L., Y.Y, and Y.Z. designed the study. Q.X. and T.L. performed the benchmarking analyses. Z.L. and C.L. contributed to the literature review and public data collection. Q.X. and Y.Z. drafted the manuscript and all authors have reviewed and commented the final manuscript.

---

**Figure S1:**
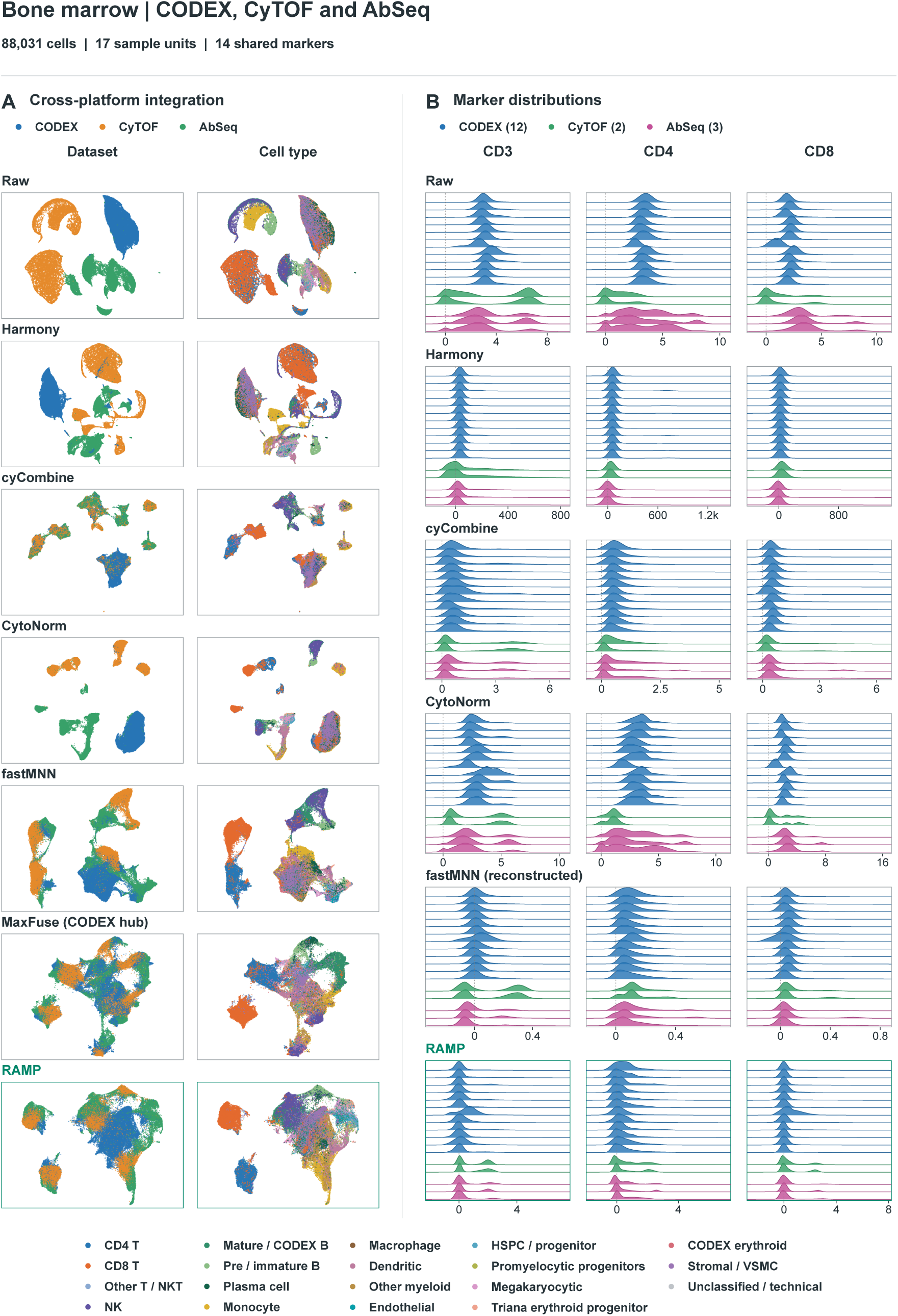
Bone marrow benchmark. CODEX, CyTOF and Triana AbSeq: 88,031 cells, 17 sample units and 14 shared markers. (A) All frozen cells and the original display labels are retained. (B) CD3, CD4 and CD8 distributions for all 17 units (12 CODEX, two CyTOF and three AbSeq). Raw values are displayed after arcsinh transformation with cofactor 1. This benchmark is distinct from the CD19-recovery application (Fig. 4).

**Figure S2:**
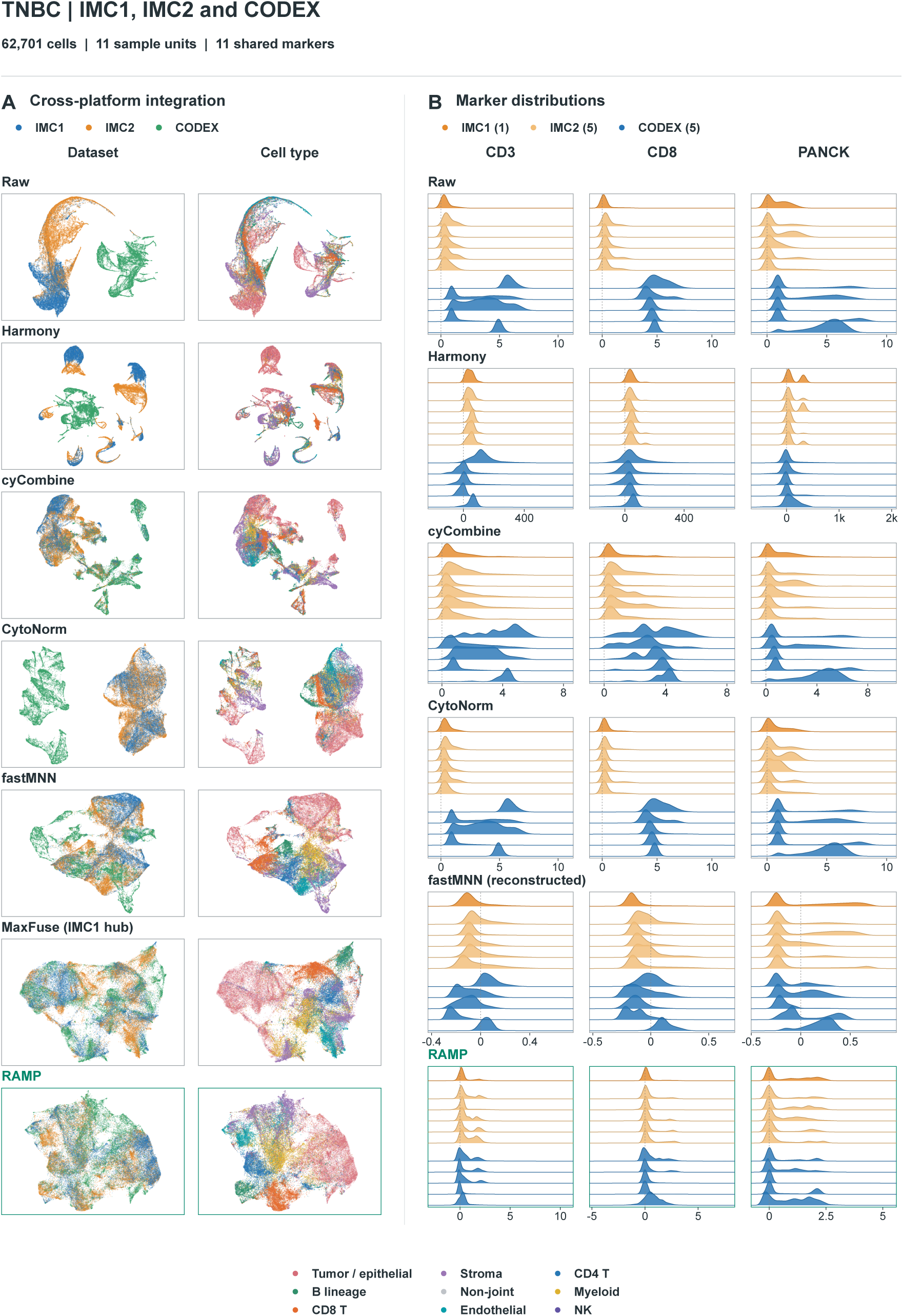
TNBC benchmark. Tietscher IMC (IMC1), Meyer IMC (IMC2) and Shiao CODEX: 62,701 cells, 11 sample units and 11 shared features. (A) All frozen cells and the original display labels are retained. (B) CD3, CD8 and PANCK distributions for all 11 units (one IMC1, five IMC2 and five CODEX). Raw values are displayed after arcsinh transformation with cofactor 1; IMC1 and IMC2 use different orange shades.

**Figure S3:**
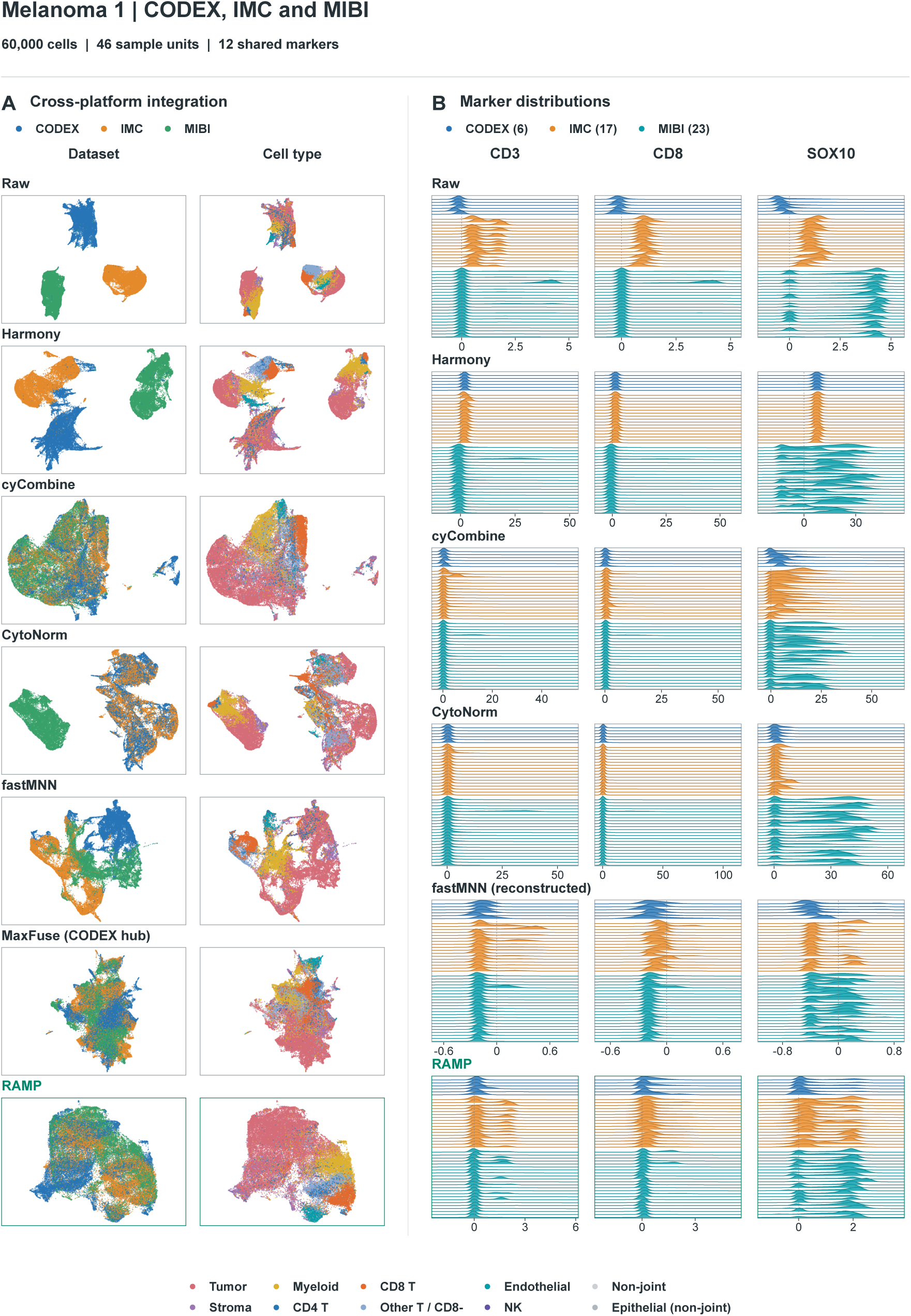
Three-platform melanoma benchmark. Hickey CODEX, Hoch IMC and Ptacek MIBI: 60,000 cells, 46 sample units and 12 shared features. (A) All frozen cells are displayed. (B) CD3, CD8 and SOX10 distributions for all 46 units (six CODEX, 17 IMC and 23 MIBI). Raw values are displayed after arcsinh transformation with cofactor 1. The Hickey SOX10/S100 channel serves as the corresponding SOX10 feature.

**Figure S4:**
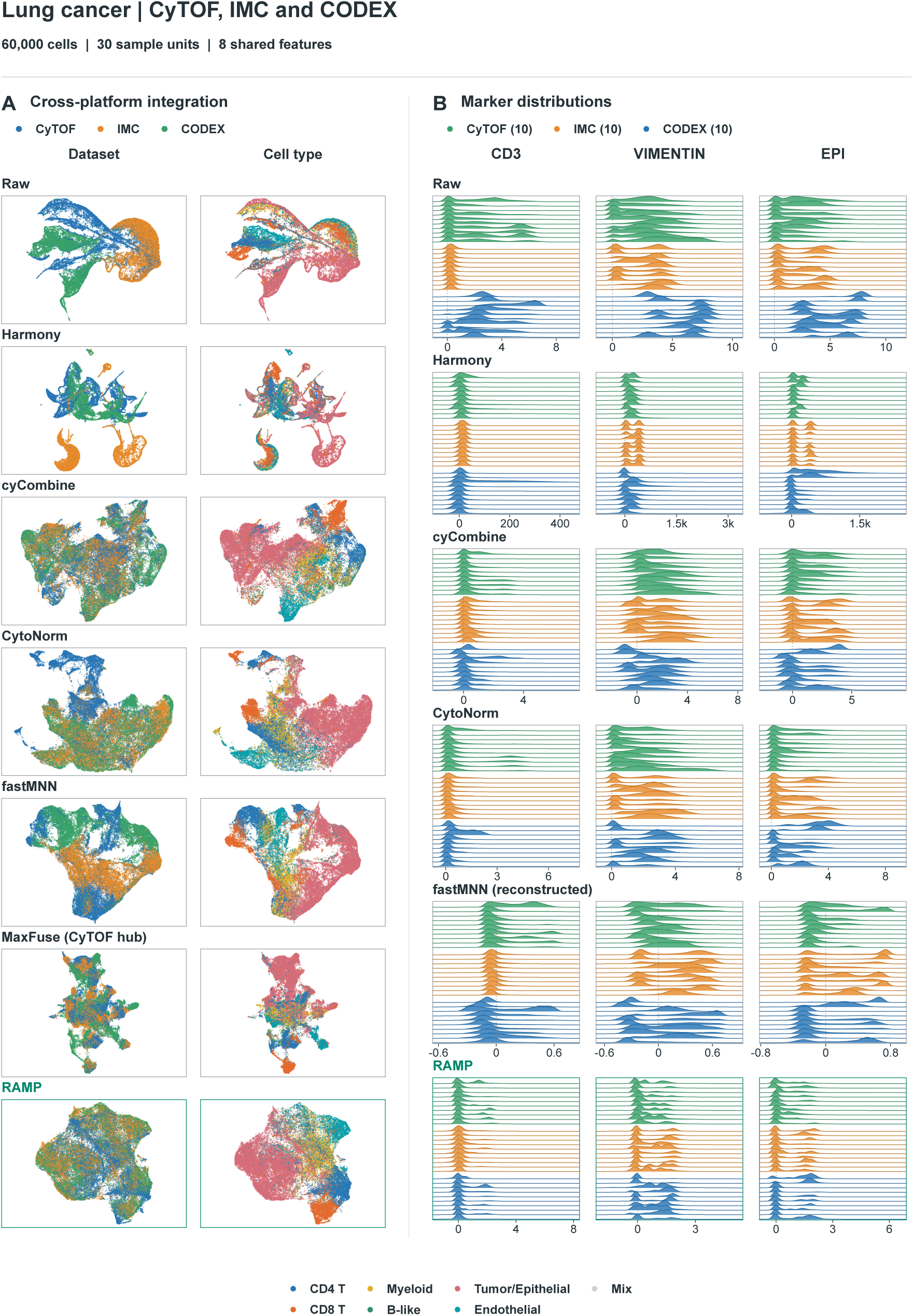
Original lung cancer benchmark. Senosain CyTOF, Cords IMC and Monkman CODEX: 60,000 cells, 30 sample units and eight shared features. (A) All frozen cells are displayed. (B) CD3, vimentin and the corresponding epithelial-marker feature (EPI) for all 30 units, ten per platform. Raw values are displayed after arcsinh transformation with cofactor 1. EPI denotes corresponding platform-specific channels rather than identical antibody measurements. This task is distinct from the lung HLA-ABC application (Supplementary Fig. S5).

**Figure S5:**
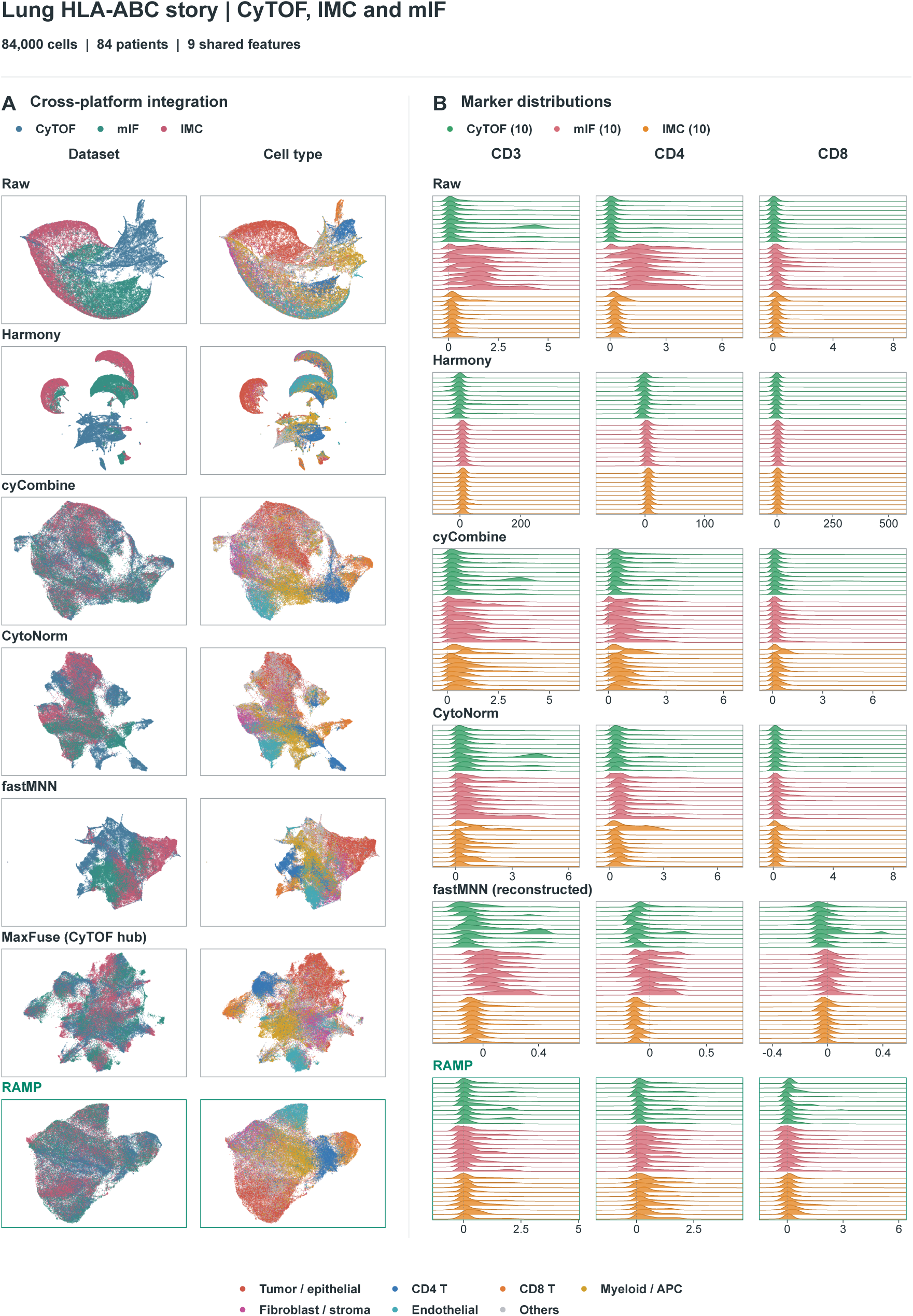
Lung HLA-ABC application cohort. Senosain CyTOF, Cords IMC and Zheng mIF: 84,000 reference cells from 84 patients and nine corresponding shared features. (A) All reference cells are displayed using the current story UMAP protocol. (B) CD3, CD4 and CD8 for a fixed subset of ten patients per platform (30,000 cells). Patients were selected by deterministic ID-hash ordering, independently of expression or performance; each selected patient contributes all 1,000 frozen reference cells. Raw display transformations are arcsinh with cofactor 5 for CyTOF and cofactor 1 for IMC/mIF.

**Figure S6:**
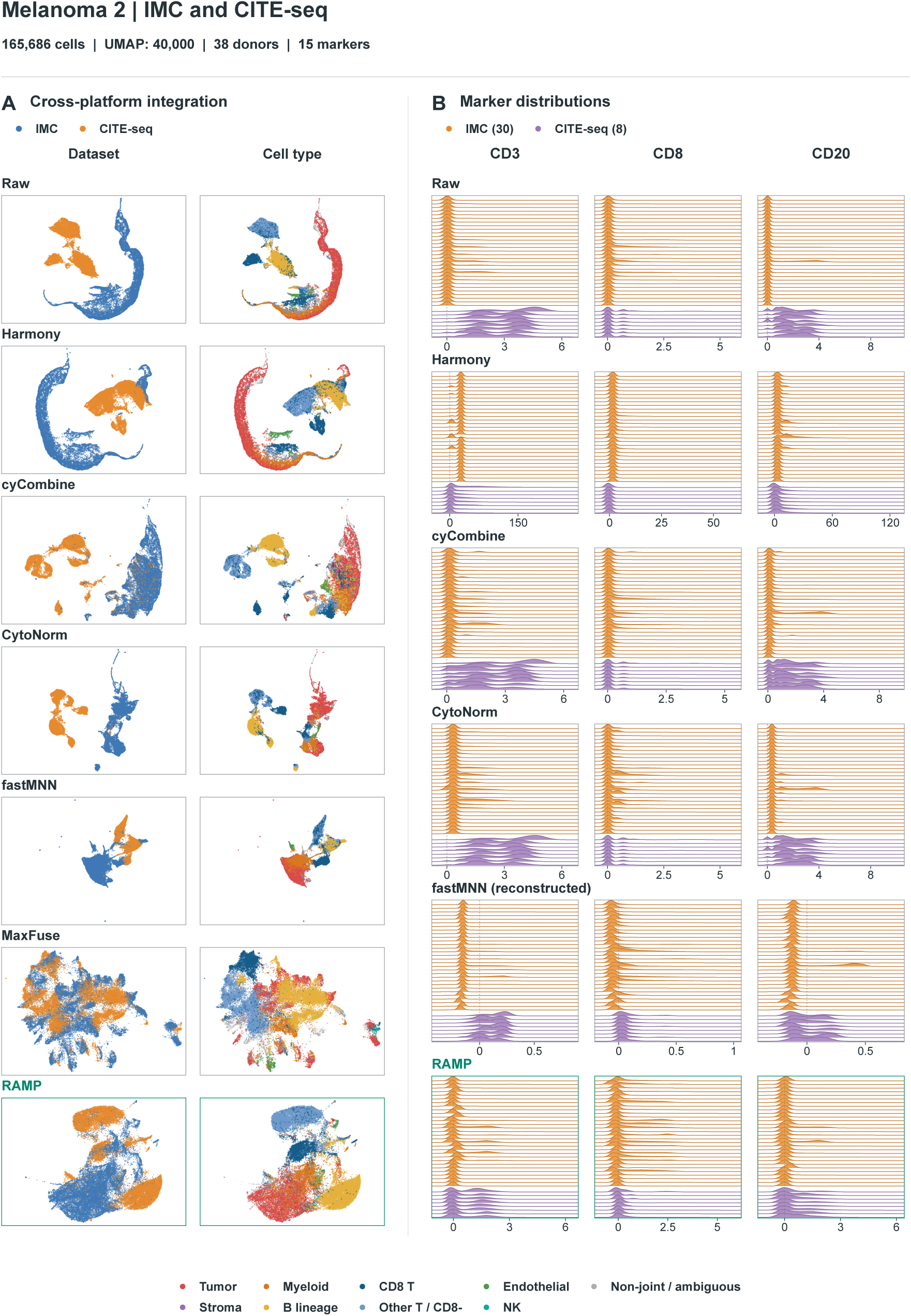
Melanoma IMC and CITE-seq application cohort. Moldoveanu IMC and Engelbrecht CITE-seq: 165,686 cells, 38 donors and 15 shared proteins. (A) All methods display the same 40,000 cells used in the story, 20,000 per platform, with approximately balanced donor contributions within each platform. (B) CD3, CD8 and CD20 distributions retain all 165,686 frozen cells and all 38 donors (30 IMC and eight CITE-seq). Raw display transformations are arcsinh with cofactor 1 for IMC and log1p for CITE-seq protein counts.

**Figure S7:**
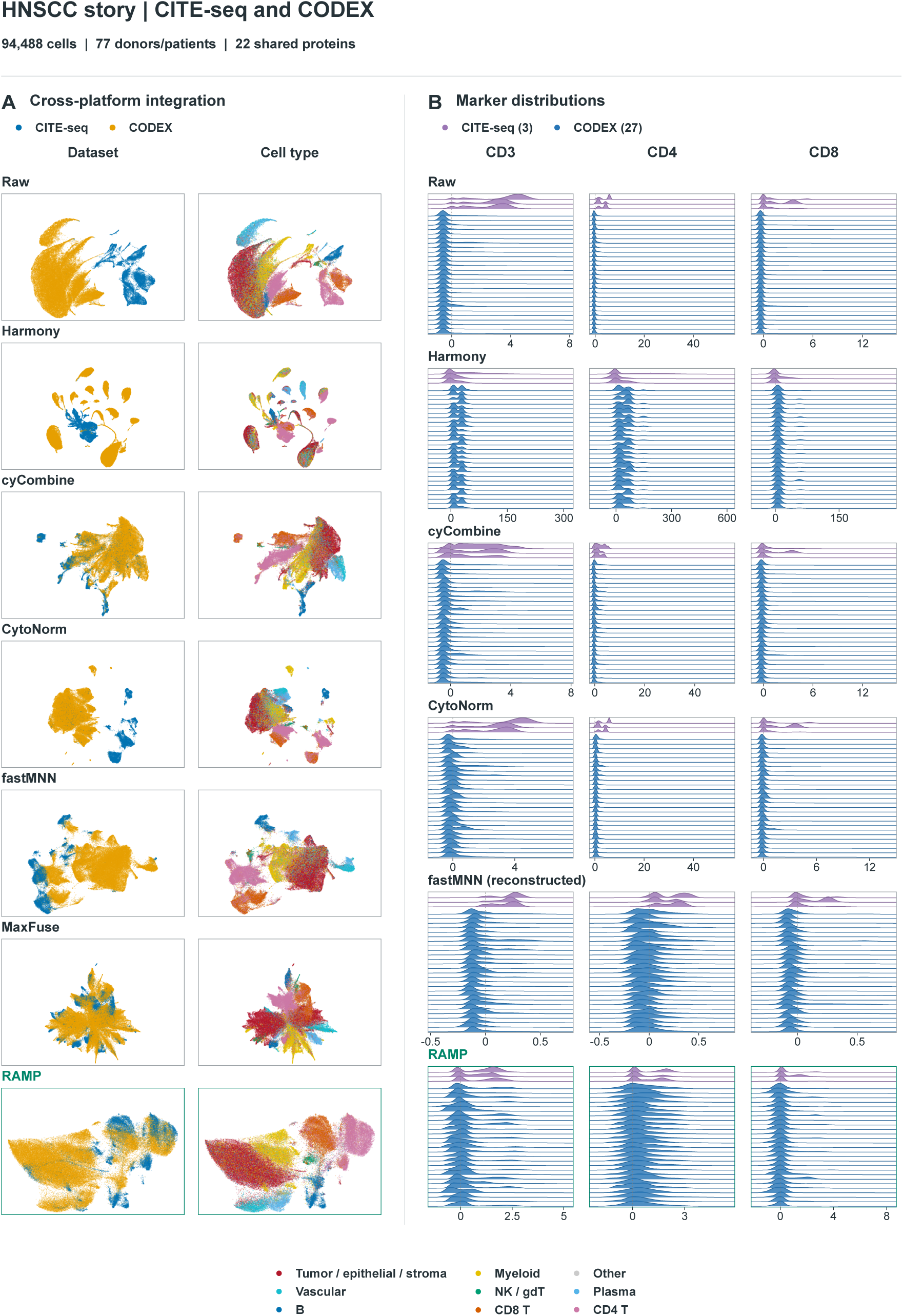
HNSCC application cohort. Rahim CITE-seq and Haist CODEX: 94,488 cells, 77 donor/patient units and 22 shared proteins. (A) All frozen cells are displayed using the story UMAP protocol. (B) CD3, CD4 and CD8 for all three CITE-seq donors and a fixed subset of 27 of 74 CODEX patients (47,488 cells). CODEX patients were selected by deterministic ID-hash ordering, independently of expression or performance; all frozen cells from selected units are retained. Raw ridges use log1p CITE-seq protein counts and provider-supplied signed CODEX values without an additional transformation.

**Figure S8:**
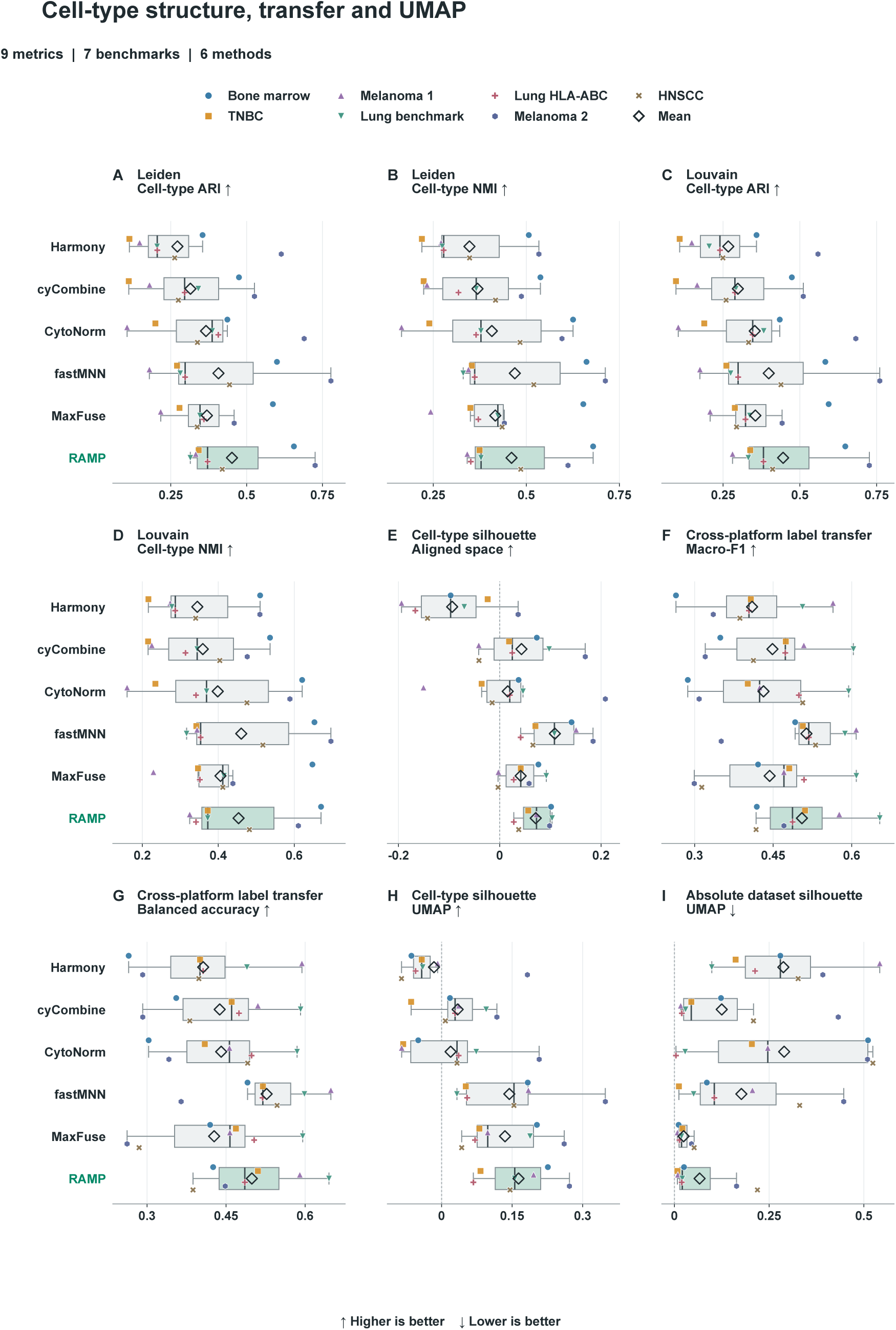
Cell-type structure, cross-platform label transfer and UMAP metrics. Nine metrics from the 23-metric reporting set are shown in their original units (Fig. 2). (A-E) Cell-type structure in the aligned representation. (F,G) Cross-platform label-transfer macro-F1 and balanced accuracy. (H,I) Cell-type silhouette and absolute dataset silhouette in UMAP. Each colored point represents one of seven benchmark scores, averaged over the 20 matched evaluation resamples. Boxes show the median and interquartile range across tasks; whiskers extend to the most extreme values within 1.5 interquartile ranges. All seven points are shown, including values beyond the whiskers. Open diamonds indicate benchmark-equal means, and arrows indicate the preferred score direction. Methods use matched evaluation cells within each task; tasks can share source cohorts.

**Figure S9:**
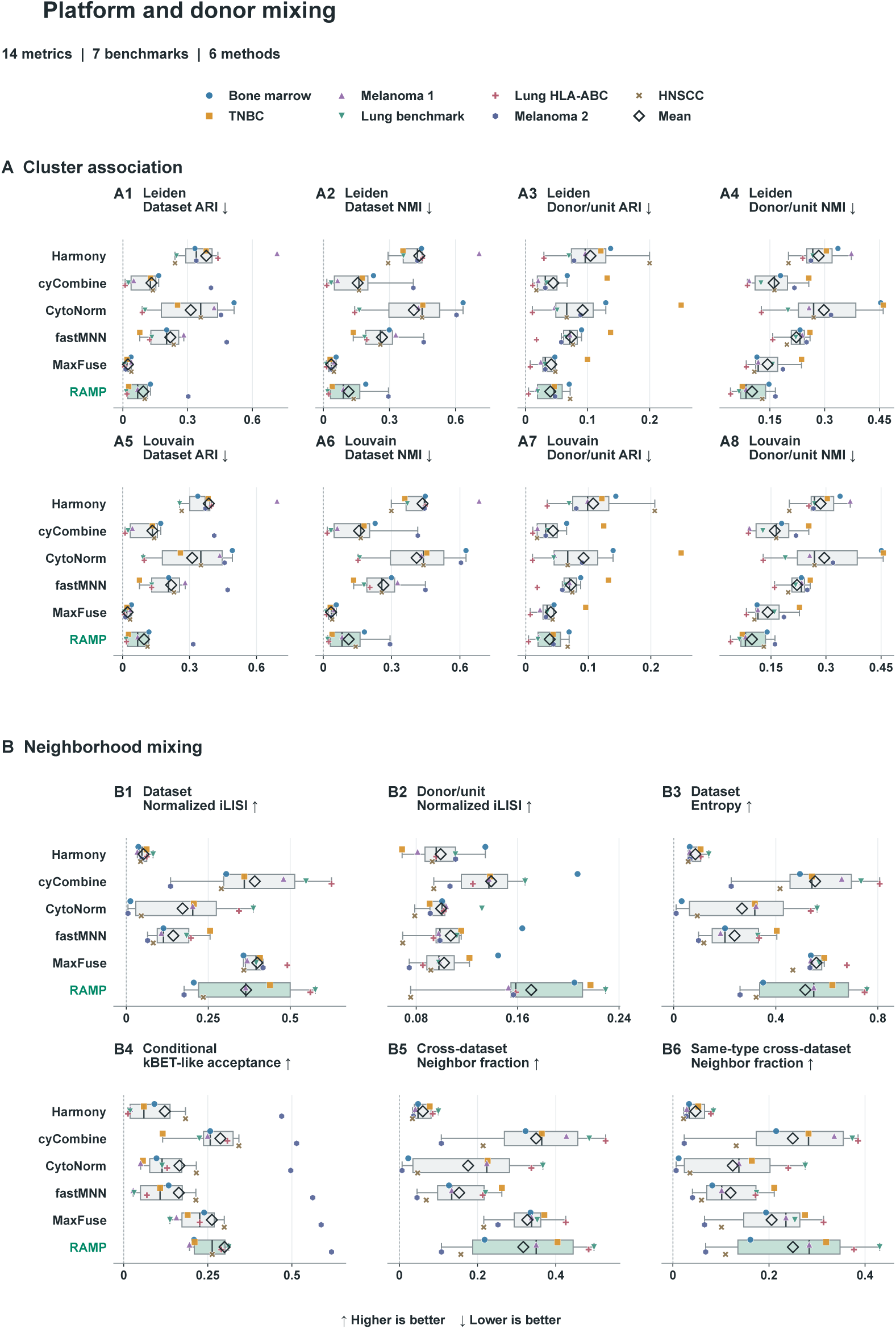
Platform and donor mixing metrics. All 14 mixing metrics from the primary reporting set are shown (Fig. 2). (A1-A8) Leiden and Louvain cluster association with dataset and sample-unit labels, measured by ARI and NMI. (B1-B6) Normalized dataset and sample-unit iLISI, dataset entropy, cell-type-conditional kBET-like acceptance, and overall and same-cell-type cross-dataset neighbor fractions. Points, boxes, diamonds and arrows use the same conventions as the structure metrics (Supplementary Fig. S8). Sample units follow the benchmark metadata and can represent donors or acquisition units. The normalized iLISI and kBET-like implementations are defined in the main Methods. The two metric figures together contain exactly the 23 primary reporting metrics (Supplementary Figs. S8 and S9); no additional metrics enter these displays.

**Figure S10:**
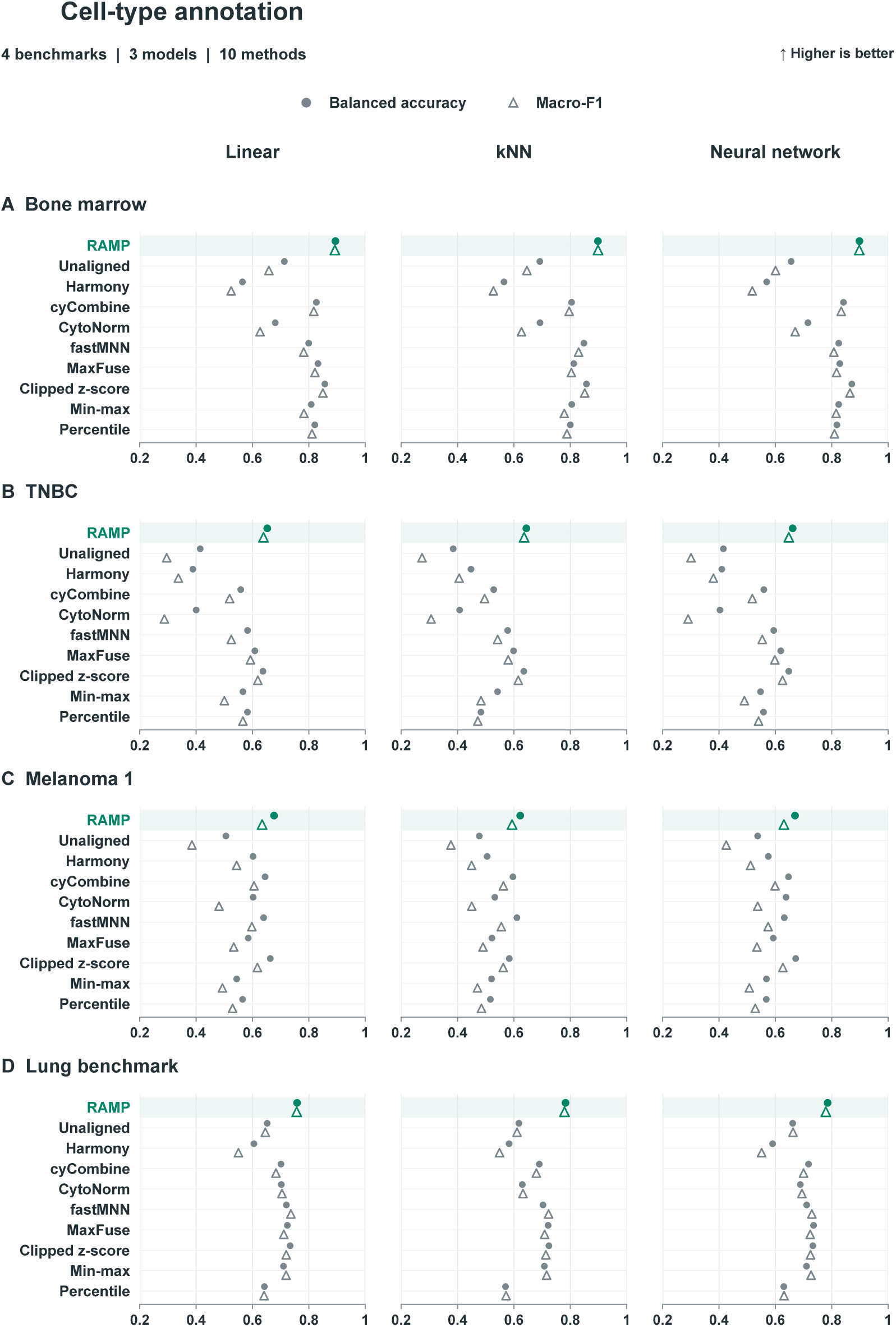
Model-specific annotation in the four original benchmarks. (A) Bone marrow, (B) TNBC, (C) original three-platform melanoma (Melanoma 1), and (D) original lung cancer benchmark. Columns show linear, k-nearest-neighbor and neural-network classifiers. Filled circles indicate balanced accuracy and open triangles indicate macro-F1; vertical offsets separate the two markers without changing their scores. Each row contains all ten processing methods. RAMP is highlighted in green; axes and method order are shared. These are the model-specific annotation components, with no averaging across models, metrics or benchmarks (Fig. 3B).

**Figure S11:**
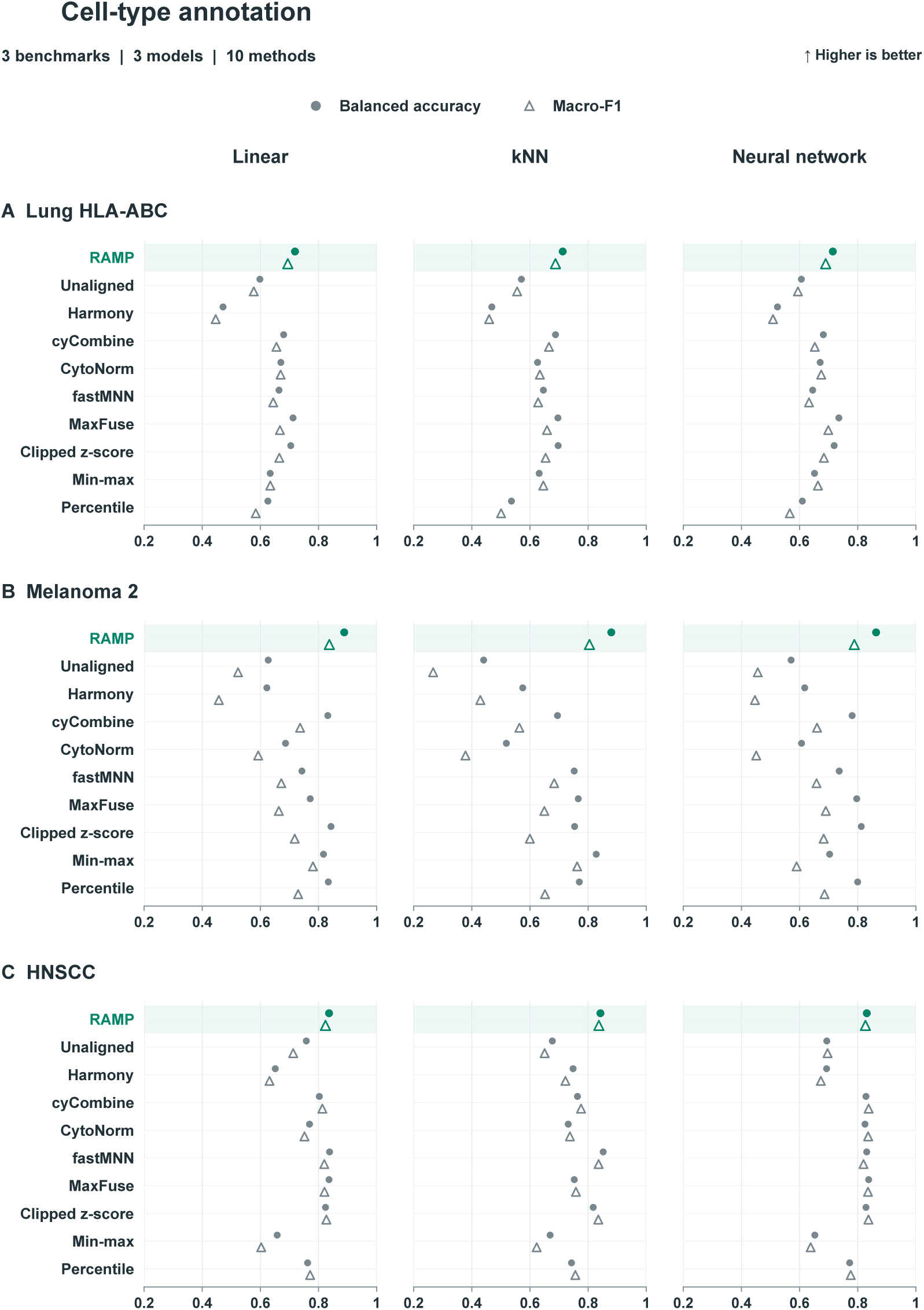
Model-specific annotation in the three application benchmarks. (A) Lung HLA-ABC, (B) melanoma IMC/CITE-seq (Melanoma 2), and (C) HNSCC. Models, symbols, axes and method order match the original-benchmark annotation plots (Supplementary Fig. S10). The Lung benchmark uses the updated Senosain reference. The two annotation figures together report all seven benchmarks (Supplementary Figs. S10 and S11), using the task-specific classification scopes described in Methods (Fig. 3B).

**Figure S12:**
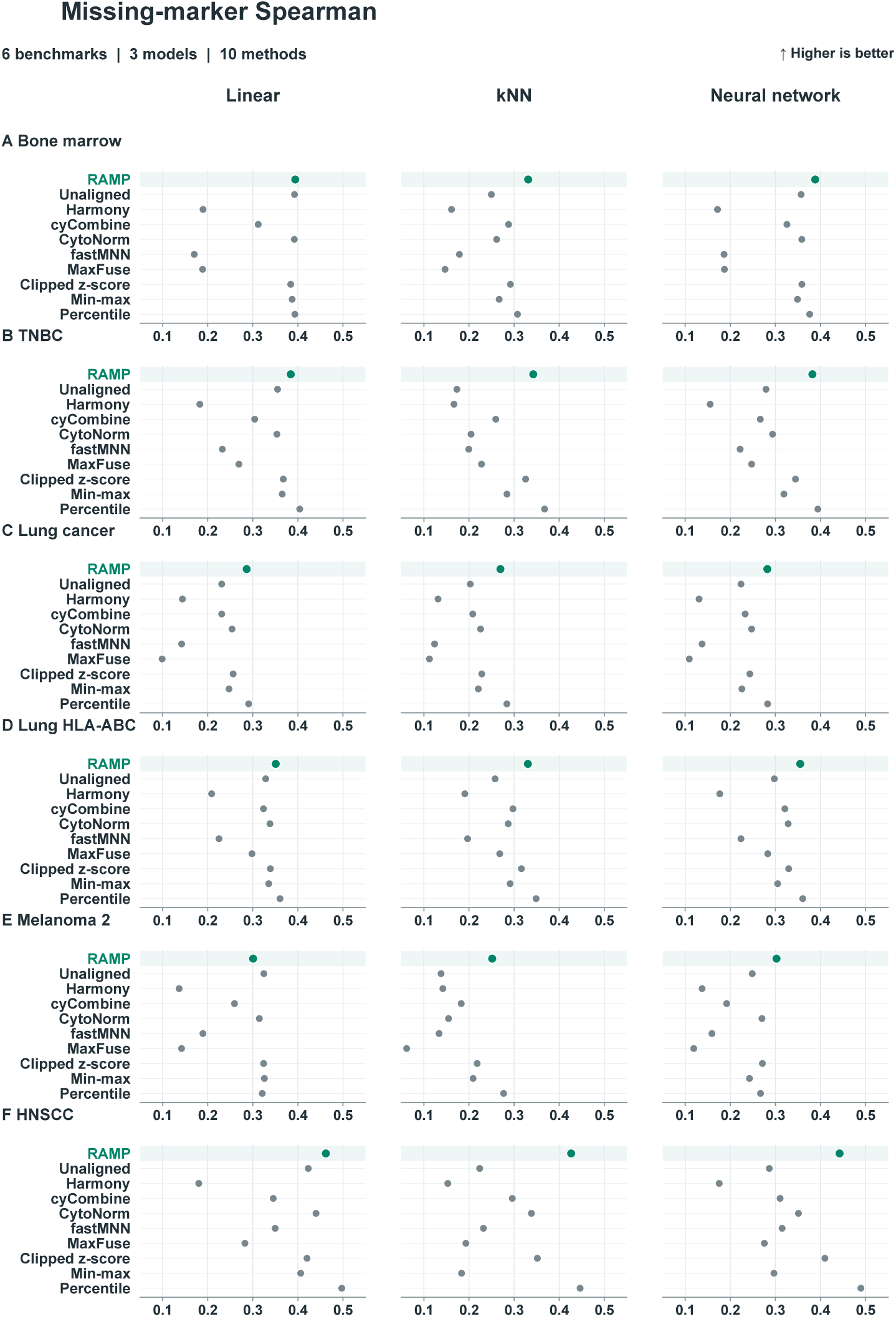
Model-specific missing-marker prediction: Spearman correlation. (A) Bone marrow, (B) TNBC, (C) original lung cancer benchmark, (D) Lung HLA-ABC, (E) melanoma IMC/CITE-seq and (F) HNSCC. Columns show linear, k-nearest-neighbor and neural-network regression. Each point is the frozen benchmark-level effective Spearman correlation for one processing method and model; higher values are better. Correlations are computed within eligible target sample units, with constant predictions assigned zero under the registered scoring rule. All six eligible benchmarks and ten methods are retained. The original three-platform melanoma benchmark was excluded from marker holdout because upstream marker dependencies in its distributed MIBI input could not be resolved.

**Figure S13:**
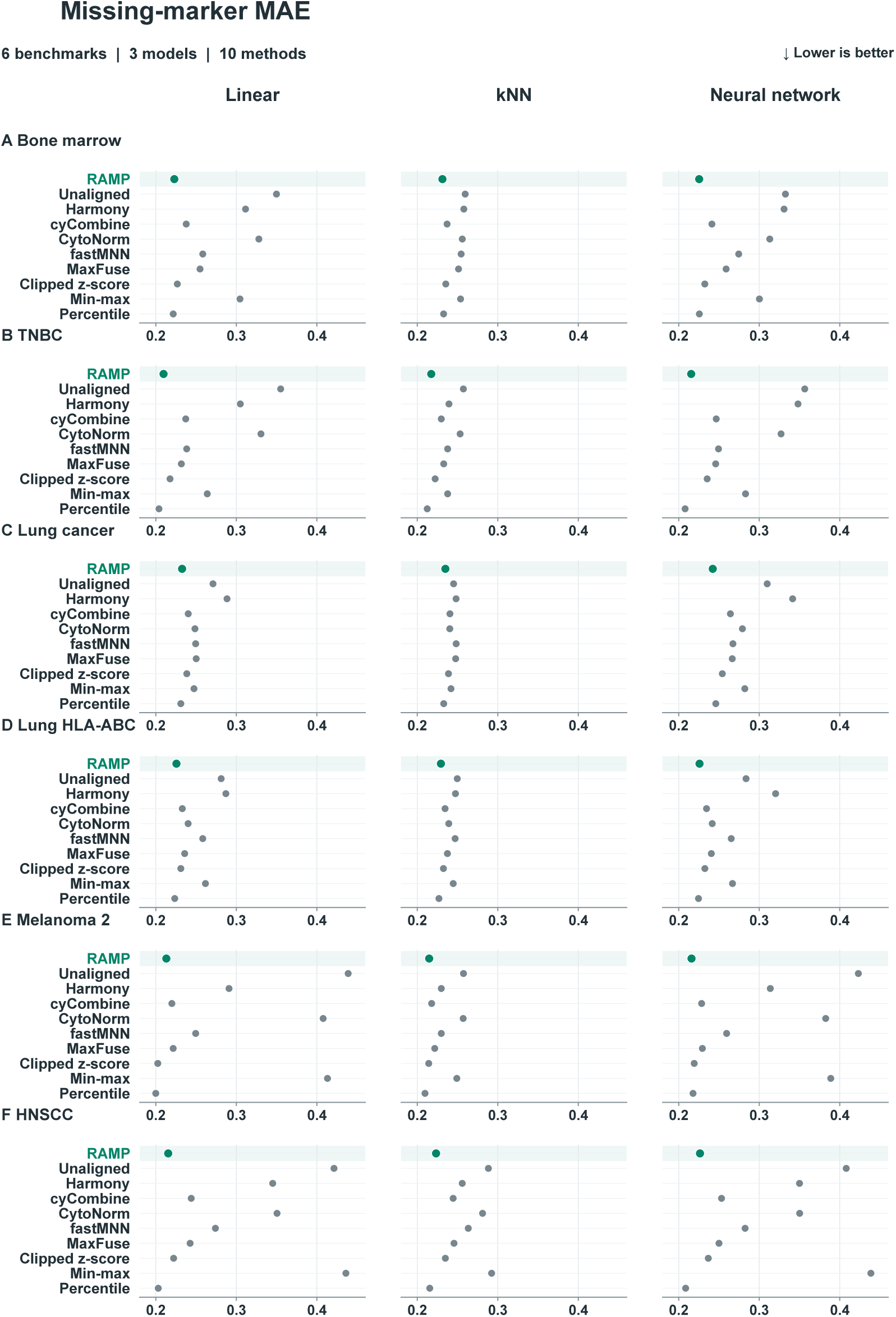
Model-specific missing-marker prediction: relative-expression MAE. Benchmark rows and model columns match the Spearman plots (Supplementary Fig. S12). Mean absolute error compares predictions clipped to [0,1] with the within-unit relative-expression target defined in Methods; lower values are better. This endpoint measures error on the common relative-expression scale, not absolute protein concentration. Axes and method order are shared across panels. The two prediction figures show the model-specific components without averaging across models or benchmarks (Fig. 3C; Supplementary Figs. S12 and S13).

